# Pathological mutations reveal the key role of the cytosolic iRhom2 N-terminus for phosphorylation-independent 14-3-3 interaction and ADAM17 binding, stability, and activity

**DOI:** 10.1101/2023.08.31.555720

**Authors:** Katharina Bläsius, Lena Ludwig, Sarah Knapp, Charlotte Flaßhove, Friederike Sonnabend, Diandra Keller, Nikola Tacken, Xintong Gao, Selcan Kahveci-Türköz, Caroline Grannemann, Aaron Babendreyer, Colin Adrain, Sebastian Huth, Jens Malte Baron, Andreas Ludwig, Stefan Düsterhöft

## Abstract

The protease ADAM17 plays an important role in inflammation and cancer and is regulated by iRhom2. Mutations in the cytosolic N-terminus of human iRhom2 cause tylosis with oesophageal cancer (TOC). In mice, partial deletion of the N-terminus results in a curly hair phenotype (cub). These pathological consequences are consistent with our findings that iRhom2 is highly expressed in keratinocytes and in oesophageal cancer. Cub and TOC are associated with hyperactivation of ADAM17-dependent EGFR signalling. However, the underlying molecular mechanisms are not understood. We have identified a non-canonical, phosphorylation-independent 14-3-3 interaction site that encompasses all known TOC mutations. Disruption of this site dysregulates ADAM17 activity. The larger cub deletion also includes the TOC site and thus also dysregulated ADAM17 activity. The cub deletion, but not the TOC mutation, also causes severe reductions in stimulated shedding, binding, and stability of ADAM17, demonstrating the presence of additional regulatory sites in the N-terminus of iRhom2. Overall, this study contrasts the TOC and cub mutations, illustrates their different molecular consequences, and reveals important key functions of the iRhom2 N-terminus in regulating ADAM17.

## Introduction

Oesophageal cancer (OC) is a critical global health concern, given its alarming prognosis and high mortality rate, ranking as the sixth leading cause of cancer-related death worldwide. Oesophageal squamous cell carcinoma (OSCC) and oesophageal adenocarcinoma (OAC) are the primary histological subtypes of OC. Despite their similarities, OSCC and OAC differ both epidemiologically and biologically (Rizza et al., 2015, Smyth et al., 2017, Uhlenhopp et al., 2020). The prognosis for OSCC, which comprises most OC cases, is exceptionally poor (Uhlenhopp et al., 2020, Yang et al., 2022). Notably, smoking and alcohol consumption serve as significant risk factors for OSCC development. However, the underlying molecular and genetic mechanisms driving OC remain poorly understood.

Howel-Evans syndrome, also called tylosis with oesophageal carcinoma (TOC), directly correlates with an increased risk to develop OSCC (Howel-Evans et al., 1958, Blaydon et al., 2012). This rare genetic disorder affects the seven-transmembrane spanning pseudoprotease iRhom2 (Fig. 1A) and its functions, altering EGF receptor (EGFR) signalling pathways by aggravating iRhom2-dependent ectodomain shedding (Blaydon et al., 2012).

**Figure 1:**
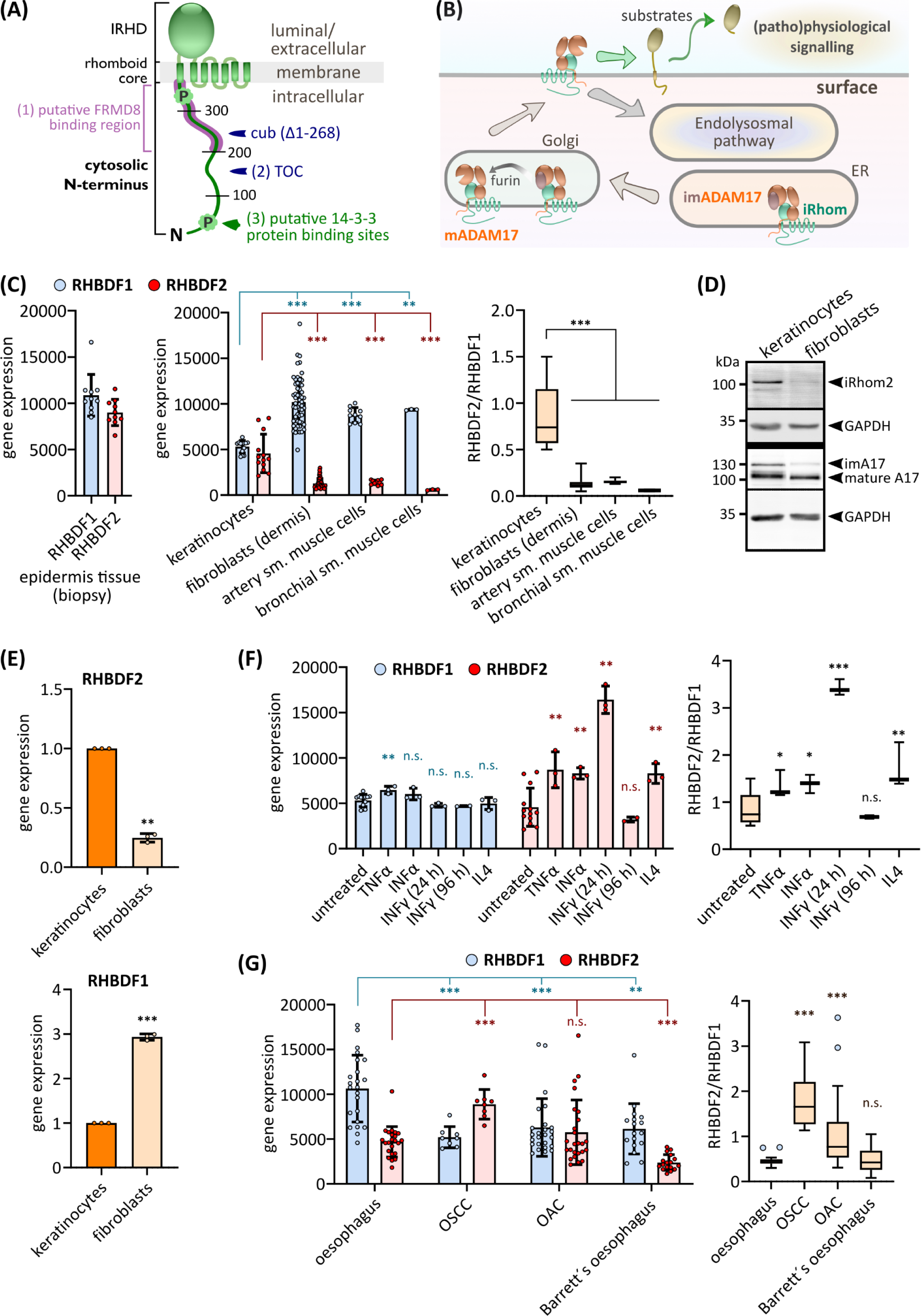
iRhom2 gene expression is high in skin and OSCC. **(A)** Structurally, iRhoms have a distinct topology composed of a rhomboid core with seven transmembrane helices (TMH). The extracellular iRhom homology domain (IRHD) is situated between TMH1 and TMH2. Additionally, iRhoms possess a large cytosolic N-terminal tail: 375 residues in murine iRhom2 and 403 residues in human iRhom2. Relative positions of key characteristics are labelled for murine iRhom2: (1) putative FRMD8 binding site; (2) TOC mutation site - in human iRhom2: I186T (murine position: 156)(Blaydon et al., 2012), D188N (murine position: 158)(Blaydon et al., 2012), D188Y (murine position: 158)(Mokoena et al., 2018), P189L (murine position: 159)(Blaydon et al., 2012), R197S (murine position: 167)(Qu et al., 2019); (3) phosphorylation sites and putative 14-3-3 binding sites. **(B)** Role of iRhoms in ADAM17 biology: iRhoms regulate ADAM17 by binding to immature proADAM17 in the ER and transporting it to the Golgi for maturation. The inhibitory prodomain is then proteolytically removed by furin-like proteases. This process allows mature ADAM17 in complex with iRhoms to localise to the cell surface and initiate shedding of its substrates, enabling various pathways including TNFα and EGFR signalling. **(C)** Curated Affymetrix Human Genome U133 Plus 2.0 Array platform data from public repositories (Hruz et al., 2008) were assessed with the GENEVESTIGATOR software. The GENEVESTIGATOR platform allows for the comparison of mRNA expression of genes between samples of different experimental data sets: Gene expression levels of iRhom1 (RHBDF1) and iRhom2 (RHBDF2) from healthy human epidermis tissue (n = 10), untreated primary human keratinocytes from healthy tissue (n = 13), untreated primary human fibroblasts from healthy dermis tissue (n = 57), untreated primary human artery smooth muscle cells from healthy heart tissue (n = 10), untreated primary human bronchial smooth muscle cells from healthy heart tissue (n = 10) were analysed. To compare the gene expression levels of iRhom1 and iRhom2 in one sample, the iRhom2/iRhom1 ratio was calculate for each sample. **(D)** Immunoblot of cell lysates from primary keratinocytes or primary fibroblasts. n = 3. **(E)** RT-qPCR of primary keratinocytes and primary fibroblasts. n = 3. **(F-G)** Data from curated Affymetrix Human Genome U133 Plus 2.0 Array platform data as described in (C): **(F)** Gene expression levels of iRhom1 and iRhom2 in primary keratinocytes from healthy tissue treated with: untreated control (n = 13) or treated for 24 h with indicated proinflammatory stimulants (n = 3) or treated for 96 h with IFNγ (n = 2). **(G)** Gene expression levels of iRhom1 and iRhom2 in healthy human oesophagus squamous tissue (n = 23) compared to expression levels in human oesophageal squamous cell carcinoma (OSCC; n = 8), human oesophageal adenocarcinoma (OAC; n = 26) and Barrett’s oesophagus (n = 17).

Shedding, the proteolytic release of ectodomains from membrane proteins, is a crucial mechanism in many physiological and pathological processes. The released soluble ectodomains, which are often receptors or receptor ligands, exhibit potent biological activity. iRhoms, pseudoproteases of the rhomboid family, play a crucial role in the shedding process by forming a complex with the transmembrane sheddase ADAM17 (A-disintegrin and metalloproteinase 17)(Siggs et al., 2012, Christova et al., 2013, Siggs et al., 2014, Li et al., 2015).

ADAM17 has a broad spectrum of substrates, including cytokines, growth factors and cytokine receptors such as tumour necrosis factor (TNF)(Black et al., 1997), transforming growth factor α (TGFα) and amphiregulin (AREG)(Peschon et al., 1998, Sahin et al., 2004). Release of these substrates is crucially involved in various (patho-)physiological processes (Peschon et al., 1998, Blaydon et al., 2011) highlighting the central role of ADAM17 in development and in the immune response but also in chronic inflammation and cancer. Additionally, the multi-faceted role of ADAM17 in virus entry has been recently demonstrated (Lambert et al., 2005, Palau et al., 2020, Yuan et al., 2021, Jocher et al., 2022, Niehues et al., 2022, Zaruba et al., 2022).

The formation of the iRhom-ADAM17 complex is a necessary step for ADAM17 maturation (Fig. 1A,B)(Christova et al., 2013, Maretzky et al., 2013, Grieve et al., 2017, Düsterhöft et al., 2021, Kahveci-Türköz et al., 2023). On the cell surface the shedding activity of mature ADAM17 is further regulated by iRhoms (Cavadas et al., 2017, Grieve et al., 2017). In mammals, two genes for iRhoms exist: RHBDF1 (iRhom1) and RHBDF2 (iRhom2). While iRhom1 is present in most cell types, iRhom2 initially appeared to be immune cell specific, but is also upregulated during inflammatory episodes in other cell lineages such as endothelial and epithelial cells (Lu et al., 2017, Babendreyer et al., 2020, Giese et al., 2021). Consistent with this, iRhom2 deficiency in mice and humans does not cause obvious developmental defects, but impairs the immune response (Adrain et al., 2012, McIlwain et al., 2012, Kubo et al., 2022).

iRhoms contain a large N-terminal cytosolic tail that has not yet been fully characterised (Fig. 1A). The genetic disorder causing TOC is caused by single point mutations in the N-terminus (Howel-Evans et al., 1958, Ellis et al., 1994, Blaydon et al., 2012, Saarinen et al., 2012, Ellis et al., 2015, Mokoena et al., 2018). This disorder is characterised by symptoms such as leukoplakia and hyperkeratosis and increases the risk of OSCC (Chao-Chu et al., 2021). The mutations are classified as “gain of function” because they apparently upregulate ADAM17 activity, resulting in higher levels of EGFR ligands, EGFR hyperactivation in keratinocytes and increased release of TNF receptor 1, leading to reduced apoptosis (Blaydon et al., 2012, Maney et al., 2015, Hosur et al., 2018b).

In mice, a spontaneous mutation called cub (curly-bare) results in a deletion of 268 amino acid residues of the N-terminus including the TOC mutation positions, causing aberrant hair growth, hyperkeratosis and hyperproliferative wound healing (Johnson et al., 2003, Siggs et al., 2014, Hosur et al., 2018a). This cub deletion was linked to higher ADAM17-mediated shedding of AREG. However, different studies contested the notion that the cub mutation results in higher ADAM17 activity at least in *in vitro* settings (Siggs et al., 2014, Hosur et al., 2018a). Interestingly, mice carrying TOC mutations exhibit the same skin-related phenotype as cub mice (Hosur et al., 2017, Hosur et al., 2018a, Hosur et al., 2018b). Noteworthy, the deficiency in iRhom2 in mice seems to cause a thinning of epidermis in the footpad (Maruthappu et al., 2017). The phenotype of both, cub and TOC mice, can be rescued by AREG deficiency, suggesting a similar underlying process with increased AREG-dependent signalling (Hosur et al., 2018b). However, the molecular mechanism for these mutations remains unknown.

Only a few regulatory regions have been identified within the N-terminus of iRhom2, including phosphorylation sites that facilitate phosphorylation-dependent binding of members of the 14-3-3 protein family (Cavadas et al., 2017, Grieve et al., 2017). Deletion of the entire N-terminus appears to destabilise the iRhom-ADAM17 complex, resulting in loss of mature ADAM17. The adapter protein FRMD8, also known as iTAP (iRhom tail-associated protein) has been identified as an essential binding partner of the N-terminus of iRhom, which has a stabilising function for iRhom and thus for the iRhom-ADAM17 complex (Künzel et al., 2018, Oikonomidi et al., 2018). This seems to explain the loss of mature ADAM17 when the N-terminus is missing. Notably, part of this FRMD8-binding region is lost in the cub deletion, while the TOC mutations lie outside this region. Recently, it was shown that the N-terminus is cleaved off by a signal peptidase complex and migrates to the nucleus, where it influences gene expression (Dulloo et al., 2022, Zanotti et al., 2022).

In this study, we observed high expression levels of iRhom2 in skin tissue samples and primary keratinocytes under physiological conditions, providing an explanation for the distinct skin-related phenotype in humans and mice with TOC or cub, respectively. Notably, iRhom2 and ADAM17 expression levels were significantly upregulated in OSCC patient samples, which is consistent with the higher risk for OSCC in TOC patients.

It is important to separate the direct molecular effects of both mutation types (single point mutation vs large deletion) on the functions of the iRhom2-ADAM17 complex from downstream effects: *In vivo*, altered ADAM17 activity and the mutated soluble iRhom2 N-terminus in the nucleus will have long-term effects on gene expression as well as ADAM17-dependent signalling pathways. Here, we focused on the direct molecular consequences of the TOC single point mutations and the large cub deletion. We identified a highly conserved motif that contains all known TOC sites and represents a phosphorylation-independent 14-3-3 protein interaction site. Disruption of this site results in loss of regulation of ADAM17 activity, leading to increased constitutive shedding. In contrast, the pathological murine cub deletion covers a larger part of the N-terminus, including the TOC site. Therefore, the cub deletion also causes higher constitutive shedding activity of ADAM17. However, the cub deletion loses the ability to promote stimulated/activatable ADAM17 activity. In addition, we found that this deletion decreases binding to ADAM17 and causes accelerated degradation of ADAM17.

Overall, our results provide deeper insights into the molecular mechanisms controlling the iRhom2-ADAM17 complex through the N-terminus of iRhom2 by demonstrating a distinct effect of TOC site disruption, in contrast to the multiple overlapping and therefore more severe effects of the cub deletion.

## Results

### iRhom2 expression levels are high in skin and OSCC

The observed phenotypes of iRhom2 deficiency and cub deletion, as well as the pathologies associated with the human TOC mutations, clearly point to a central role of iRhom2 in epidermal tissue. While high iRhom2 expression is often primarily associated with immune cells, its expression pattern in the epidermis is not yet well defined. Hence, we analysed gene expression data from skin biopsies of healthy individuals obtained from publicly available microarray data (Hruz et al., 2008). We indeed observed a robust expression of iRhom2 in the skin samples (Fig. 1C). Moreover, primary human keratinocytes have significantly higher levels of iRhom2 expression compared to primary fibroblasts, artery smooth muscle cells, and bronchial smooth muscle cells (Fig. 1C). Notably, the ratio of iRhom2 to iRhom1 is substantially higher in keratinocytes than in other primary cell types (Fig. 1C; Fig. S1A), indicating a pivotal role for iRhom2 in regulating ADAM17 activity in keratinocytes *in vivo*. To verify the microarray data, we analysed iRhom1 and iRhom2 protein levels via immunoblotting and gene expression via RT-qPCR in primary keratinocytes and fibroblasts. Indeed, in agreement with the microarray data, we found higher protein levels and gene expression of iRhom2 in keratinocytes compared to fibroblasts (Fig. 1D,E). This expression pattern aligns with the predominantly skin-related manifestations of TOC and cub. We found a similar pattern in murine primary keratinocytes, again highlighting iRhom2’s importance in regulating ADAM17 activity in the skin (Fig. S1B).

Considering the known induction of iRhom2 expression in response to proinflammatory stimuli (Babendreyer et al., 2020, Giese et al., 2021), we investigated whether the observed iRhom2 expression patterns in primary keratinocytes were due to pre-activation by proinflammatory conditions. We found a significant increase in iRhom2 gene expression in primary keratinocytes 24 hours after treatment with proinflammatory stimuli, while iRhom1 showed no such elevation (Fig. 1F). This finding supports that the high iRhom2 expression in primary keratinocytes is likely the basal level under physiological conditions which can be further increased under inflammatory conditions. Furthermore, the gene expression pattern of untreated keratinocytes is also consistent with the basal iRhom2 and iRhom1 gene expression pattern in healthy human skin biopsies (Fig. 1C).

Since TOC mutations also pose an increased risk of developing OSCC, we examined iRhom2 expression in OSCC samples and compared them to healthy oesophagus squamous tissue, as well as samples from OAC and a premalignant condition, called Barrett’s oesophagus. In OSCC samples, we observed a significant upregulation of iRhom2, accompanied by a downregulation of iRhom1, compared to healthy oesophagus squamous tissue (Fig. 1G). The ratio of iRhom2 to iRhom1 was four times higher in OSCC samples than in the healthy tissue. While OAC samples also showed a significantly elevated iRhom2-iRhom1 ratio, the increase was less pronounced (Fig. 1G). On the other hand, samples from Barrett’s oesophagus exhibited no change in the ratio but displayed an overall reduction in iRhom1 and iRhom2 gene expression. Furthermore, we found a significant upregulation of ADAM17 gene expression exclusively in OSCC samples (Fig. S1C). In contrast, ADAM10, a close relative of ADAM17 with partially overlapping substrate spectrum and independent of iRhom functions, did not show this expression pattern (Fig. S1C). The gene expression of EGFR ligands, TGFα and AREG, crucial ADAM17 substrates, did not show significant changes in OSCC compared to healthy tissue (Fig. S1C). To ensure that the elevated levels of iRhom2 in OSCC were not due to persistent proinflammatory conditions or the presence of tumour microenvironment-associated immune cells, we examined proinflammatory markers and immune cell markers. We observed no significantly elevated gene expression of TNF or IL6 (Fig. S1C), and only minimal gene expression of CD14, suggesting the presence of some monocytes/macrophages in the samples but not in sufficient quantities to explain the high iRhom2 levels in OSCC (Fig. S1D; Fig. S2A,B).

In summary, the gene expression patterns of iRhom2 and ADAM17 support the notion of an important role for iRhom2-dependent ADAM17 activity in both skin and OSCC, which is consistent with the pathological consequences of TOC single point mutations and cub deletion.

### The N-terminal tail represents a large intrinsically disordered part of iRhom2

To investigate the molecular mechanisms underlying the functions of the N-terminal tail of iRhom2, we performed *in silico* structural studies. Our bioinformatic analyses predict that the N-terminal tail of iRhom2 has a largely unstructured/intrinsically disordered nature (Fig. 2A), which is typical for protein-protein interaction interfaces (Dyson, 2016, Arakawa et al., 2023). Using the A.I.-based AlphaFold2 algorithm, we predicted the structural makeup of the N-terminus, which revealed an overall low pLDDT and PAE, again confirming a lack of higher structural regions (Fig. 2B-D). Some isolated secondary structures were predicted, but the N-terminus seems to be primarily intrinsically disordered (Fig. 2A-F), which is a hallmark to promote protein-protein interaction.

**Figure 2:**
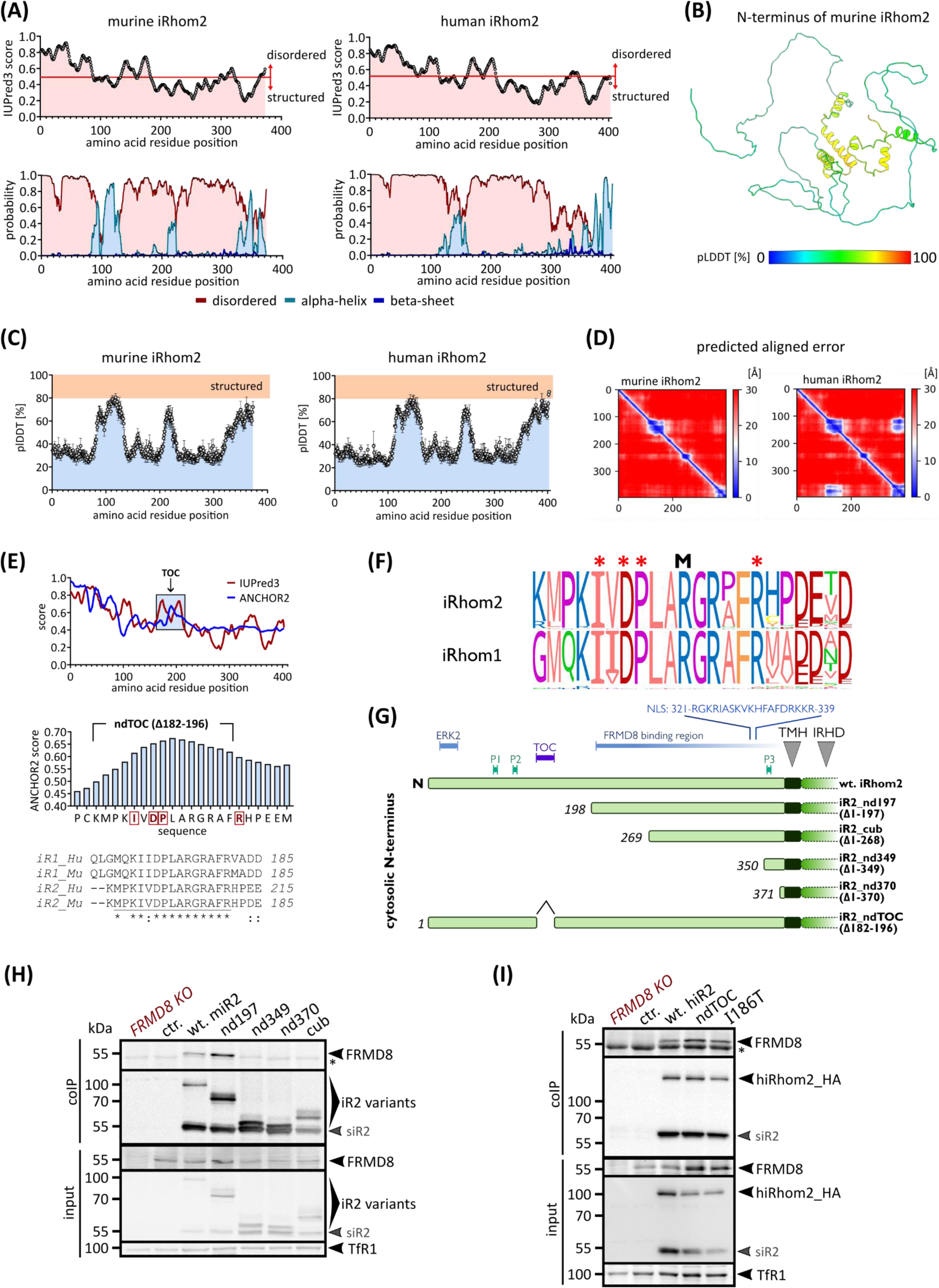
The cytosolic N-terminal tail of iRhom2 is an intrinsic disordered hub for regulation and differentially controls levels of mature ADAM17. **(A)** Using IUPred3 (Meszaros et al., 2009, Erdos et al., 2021) and NetSurfP-2.0 (Klausen et al., 2019), we assessed the probability of disordered regions and the presence of different structural elements in the cytosolic N-terminus of murine and human iRhom2 (residue 1 to 375). IUPred3 score describes the probability of disordered regions. NetSurfP-2.0 output gives the probability of the presence of disorder, alpha-helices, and beta-sheets. (B) *Ab initio* structural modelling of the N-terminus of murine iRhom2 using the deep learning algorithm AlphaFold2. Best ranked model is depicted in cartoon representation coloured according to pLDDT (predicted local distance difference test in [%]) score. **(C)** Consistent with (A), the structural prediction of the N-terminus of murine and human iRhom2 using AlphaFold2 demonstrates only small locally structured regions: Hence the iRhom2 N-terminus is largely disordered, as indicated by the overall low average pLDDT score per sequence position of the five predicted, ranked models. **(D)** The predicted align error (PAE) of the best ranked structural models support this finding: the individual amino acid residues in the disordered/unstructured N-terminus are largely not fixed in their position relative to each other and therefore have a high spatial uncertainty (20 - 30 Å). **(E)** Using the bioinformatic tools IUPred3 and ANCHOR2 (Mészáros et al. 2009) putative binding sites within the cytoplasmic N-terminus of human iRhom2 were identified. The plots show the calculated probability for an amino acid residue to be part of a binding site. Regions above 50% probability (ANCHOR2) and with a certain length are putative binding sites such as the highly conserved region which harbours all known TOC mutations. Our designed TOC site deletion ndTOC comprises the three main TOC mutations, but not the newest identified R197S (Qu et al., 2019). **(F)** We used the algorithm CONYAR (Kahveci-Türköz et al., 2023) to retrieve all available amino acid sequences of iRhom1 and iRhom2 from UniProtKB and compare them to identify highly conserved regions within the amino acid sequences of that gene. In the case of iRhom2 (query: RHBDF2) sequences from 381 species and in case of iRhom1 (query: RHBDF1) sequences of 301 species were extracted. The TOC site was identified by CONYAR as a highly conserved region and its conservation is shown as WebLogo representation. Asterisks indicate the positions of the natural occurring TOC mutations in human iRhom2. M indicates the position of arginine methylation found by high throughput proteomics (Hornbeck et al., 2015). **(G)** Overview of the relative position of key regulatory sites and binding interfaces in wt-iRhom2 and in our designed N-terminal deletions. Phosphorylation sites (P1, P2 and P3) are binding sites for 14-3-3 proteins. NLS: predicted nuclear localisation sequence. **(H,I)** To analyse binding between iRhom2 constructs and endogenous FRMD8, coIPs were performed using the indicated iRhom2 constructs (with HA tag) as bait. HEK293 cells stably expressing the indicated murine iRhom2 deletions (H) or human iRhom2 constructs (I) as well as GFP (ctr.) as negative control were used. At around 55 kDa the shorter fragment of cleaved iRhom2 (siR2) as reported before (Adrain et al., 2012, Christova et al., 2013) is visible. This fragment was described as consequence of cleavage by signal peptidase complex (Dulloo et al., 2022, Zanotti et al., 2022) but may additionally occur as artefact after lysis as we detect inconsistent amounts of the small fragment in different experiments. n = 4.

However, while the N-terminus seems to play a crucial role in regulating the iRhom-ADAM17 complex, only a handful of direct interactors such as FRMD8, ERK2, and 14-3-3 proteins have been identified (Fig. 1A)(Cavadas et al., 2017, Grieve et al., 2017, Künzel et al., 2018, Oikonomidi et al., 2018, Schumacher et al., 2023). Using ANCHOR2, we have identified an additional region that is highly likely to be an interface for protein-protein interactions (Fig. 2E). Strikingly, this region is highly conserved and includes all known TOC mutation positions (Fig. 2E-F). In addition, R192 in the middle of this TOC site was found to be methylated in human iRhom2 by three independent high-throughput proteomic screens (Figure 2F)(Guo et al., 2014, Hornbeck et al., 2015). This further supports the TOC site as a regulatory interaction site, as protein methylation is a hallmark of protein interaction sites.

Furthermore, our bioinformatic analysis of the N-terminus of iRhom2 also indicates the presence of a nuclear localisation sequence (NLS), which is consistent with recent observations of its translocation to the nucleus after cleavage by the signal peptidase complex (Fig. 2G)(Dulloo et al., 2022).

Overall, the N-terminal tail represents a large intrinsically disordered region that provides distinct hubs for regulation and interaction. Furthermore, all known TOC mutations are part of a predicted protein-protein interaction site.

### The TOC mutations and the cub deletion result in distinct molecular consequences for mature ADAM17

To gain a comprehensive understanding of the regulatory system acting on the iRhom2-ADAM17 complex, it is important to understand the molecular similarities and differences between the TOC single point mutations and cub deletions, as they exhibit a strikingly similar phenotype in mice (Hosur et al., 2018b). While the TOC mutations affect a distinct region, the cub deletion affects a large part of the N-terminal tail encompassing different known regulatory and interaction sites such as two phosphorylation sites, the TOC region as well as the ERK2 binding site (Fig. 2G). The exact binding site of the FRMD8 interaction is not yet known, but the approximate binding region was mapped before (Fig. 2G)(Künzel et al., 2018, Oikonomidi et al., 2018). Hence, in contrast to TOC mutations the cub deletion affects most likely more regulatory aspects. To further dissect the molecular regulations of iRhom2 and its dysregulation, we designed an additional N-terminal deletion (nd197) that still contains the FRMD8 binding region and therefore retains binding to endogenous FRMD8 (Fig. 2G,H). In contrast, we could not detect FRMD8 binding for the cub deletion. We created two additional deletions (nd349 and nd370) with almost the entire N-terminus removed (Fig. 2G,H) to compare the effect of complete loss of the N-terminal tail with the partial deletions. We also hypothesised that the TOC single point mutations only gradually affect the function of the identified TOC site (Fig. 2E). Hence, we designed the deletion ndTOC (Δ182-192) to remove most of the TOC site (Fig. 2E-F). As expected, binding to FRMD8 in iRhom2 is not altered by either the TOC point mutation (I186T) or the deletion of the TOC site (ndTOC) compared to wt-iRhom2 (Fig. 2I).

The TOC mutations are autosomal dominant and the cub deletion leads to a phenotype already in heterozygosity. To analyse the direct molecular effects of these iRhom2 variants on ADAM17 regulation in combination with wt-iRhoms, we used HEK293 cells with endogenous iRhom1 and iRhom2 (Fig. S1E) as an already established and widely used model system (Cavadas et al., 2017, Sieber et al., 2022, Kahveci-Türköz et al., 2023), with epithelial morphology (Inada et al., 2016).

As iRhom levels are a rate limiting factor for forward transport and maturation of ADAM17, increased expression of either human or murine wt-iRhom2 results in more mature ADAM17 (Fig. 3A-C), as shown before (Düsterhöft et al., 2021, Kahveci-Türköz et al., 2023). Consequently, the amount of mature ADAM17 on the cell surface is also elevated (Fig. 3D,E). In contrast, expression of the murine iRhom2 cub deletion does not promote higher levels of mature ADAM17 (Fig. 3A). Furthermore, the three deletions nd197, nd349 and nd370 showed also a significant decrease in mature ADAM17 levels compared to wt-iRhom2 (Fig. 3B). We also observed a significant reduction in total ADAM17 levels in cells expressing these deletions compared to wt-iRhom2 or the negative control (Fig. 3A,B). In line with these findings the amount of mature ADAM17 on the surface was also lower in cells expressing the deletions compared to both wt-iRhom2 and even control cells (Fig. 3D), although the deletion constructs can be detected on the cell surface similarly to wt-iRhom2 (Fig. S3B). Notably, these effects seem to be FRMD8 independent since nd197 still binds endogenous FRMD8 (Fig. 2H).

**Figure 3:**
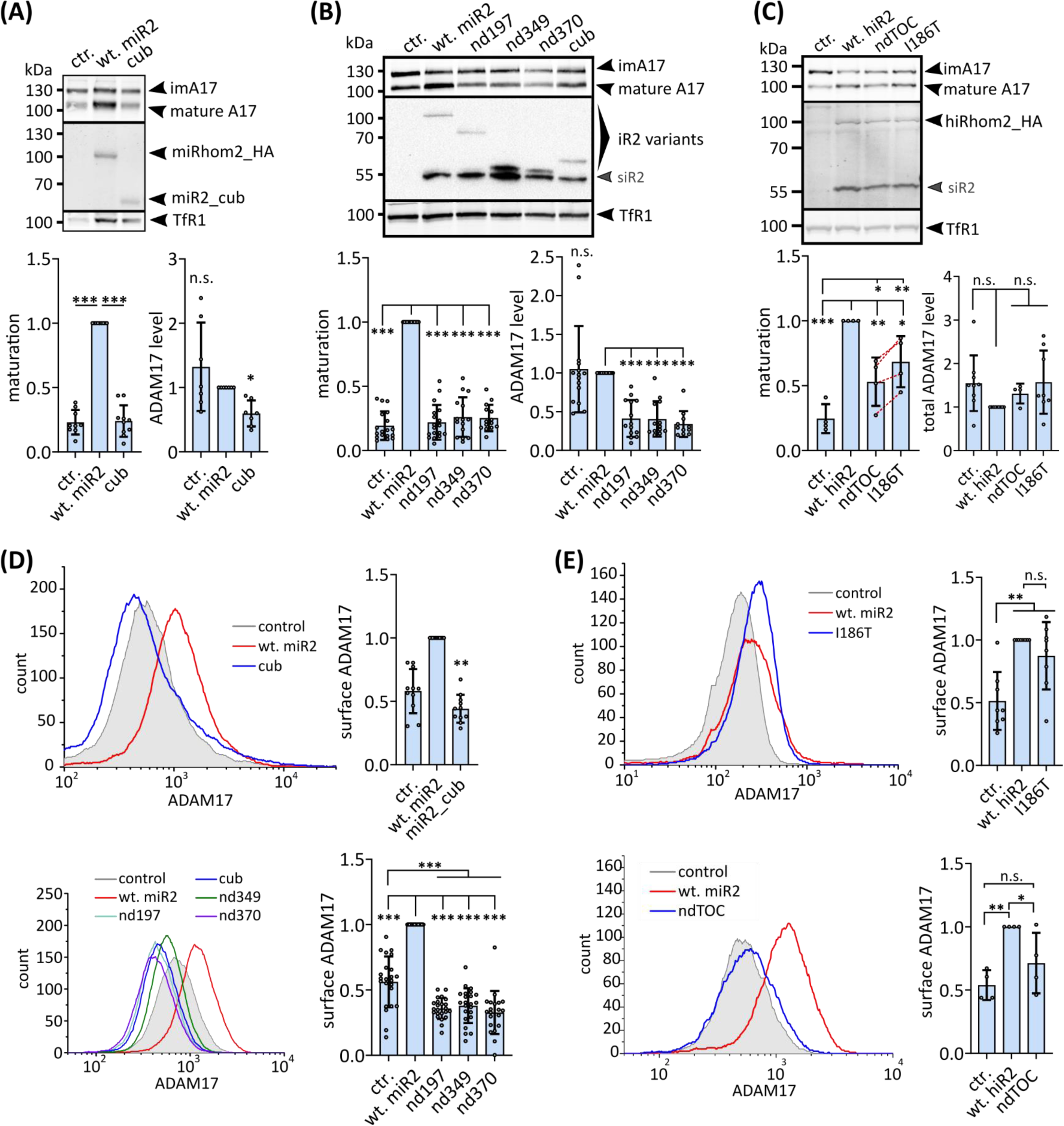
The iRhom2 N-terminus regulates levels of mature ADAM17. **(A)-(C)**Immunoblot of samples from HEK293 cells stably expressing the indicated murine iRhom2 variants (A,B) or human iRhom2 variants (C) as well as GFP (ctr.) as negative control. Glycosylated proteins from cell lysates were enriched with concanavalin A beads. HEK293 cells have endogenous iRhoms and therefore exhibit a basal level of ADAM17 maturation. The maturation level of ADAM17 can be detected by the presence of mature ADAM17 with lower molecular weight than immature ADAM17 (imA17). ADAM17 maturation was assessed by densitometric measurements and calculation of the ratio between mature ADAM17 and immature ADAM17. Total ADAM17 levels were assessed by densitometric measurements and calculation of the ratio between total ADAM17 (mature plus immature ADAM17) divided by the respective input control. Values were normalised to the respective wt-iRhom2 sample. n > 4. **(D)-(E)** Levels of surface ADAM17 in cells stably expressing indicated murine iRhom2 constructs (D) or human iRhom2 constructs (E) were measured by flow cytometry. For quantification, the geometric mean of the specific fluorescence signal was determined and normalised to the respective wt-iRhom2. n > 4.

Expression of iRhom2 with a human TOC mutation (I186T) or a deletion of the TOC site (ndTOC) also shows a reduced amount of mature ADAM17 compared to wt-iRhom2 (Fig. 3C). ndTOC seems to lead to an even stronger reduction than the single point mutation, suggesting that the complete deletion of the TOC site has a stronger effect than the single point mutation (Fig. 3C). However, in contrast to the larger deletions (nd187, cub, nd349 and nd379), we did not observe a reduction in the total amount of ADAM17 (Fig. 3C). Both iRhom2_I186T and ndTOC exhibit a cell surface presence similar to that of the wt-iRhom2 (Fig. S3C). iRhom2_I186T leads to increased ADAM17 surface localisation compared to control cells and the impact on surface ADAM17 levels is not significantly different to cells with wt-iRhom2 expression (Fig. 3E). Remarkably, ndTOC expression results in significantly lower levels of surface ADAM17 compared to wt-iRhom2 (Fig. 3E).

These findings demonstrate the importance of the cytosolic iRhom2 N-terminus in regulating ADAM17 functions. They highlight the differential effects of TOC mutations and the cub deletion on iRhom2 function and their substantial impact on ADAM17 maturation, surface localisation and overall protein levels. Overall, these findings suggest that the decrease in ADAM17 levels and downregulation of surface ADAM17 seen in the cub deletion and to a lesser extent in the TOC site mutants occur in a FRMD8-independent way. Importantly, the deletion of the TOC site in ndTOC shows a more severe molecular phenotype than the TOC single point mutations. This is consistent with single point mutations only partially disrupting TOC site functions, whereas a larger deletion has a more severe effect.

### The iRhom2 N-terminus differentially regulates ADAM17-mediated shedding

Our findings that cub and TOC mutations decreases levels of mature ADAM17 is in line with recent *in vivo* and *ex vivo* results (Siggs et al., 2014, Rabinowitsch et al., 2023). However, these findings are counterintuitive to the earlier reported elevated ADAM17 shedding activity (Brooke et al., 2014, Hosur et al., 2017, Hosur et al., 2018a, Hosur et al., 2018b). To analyse the effect of the N-terminus on ADAM17 activity, we investigated the constitutive shedding of the ADAM17 substrates TGFα, IL1R2 and AREG in HEK293 cells. As reported earlier, we found significantly increased substrate release with additional expression of wt-iRhom2 compared to control cells (Fig. 4A,B)(Düsterhöft et al., 2021, Kahveci-Türköz et al., 2023). Surprisingly, despite lower surface ADAM17 levels (Fig. 3D), ADAM17-mediated shedding was not adversely affected in cells with N-terminal deletions nd197, cub and nd349 compared to control cells (Fig. 4A,B). In fact, shedding efficiency in cells with these deletions was generally higher compared to shedding in control cells and comparable to cells expressing wt-iRhom2. The largest deletion nd370 had the lowest ADAM17-mediated shedding, but was still comparable to control cells (Fig. 4A,B).

**Figure 4:**
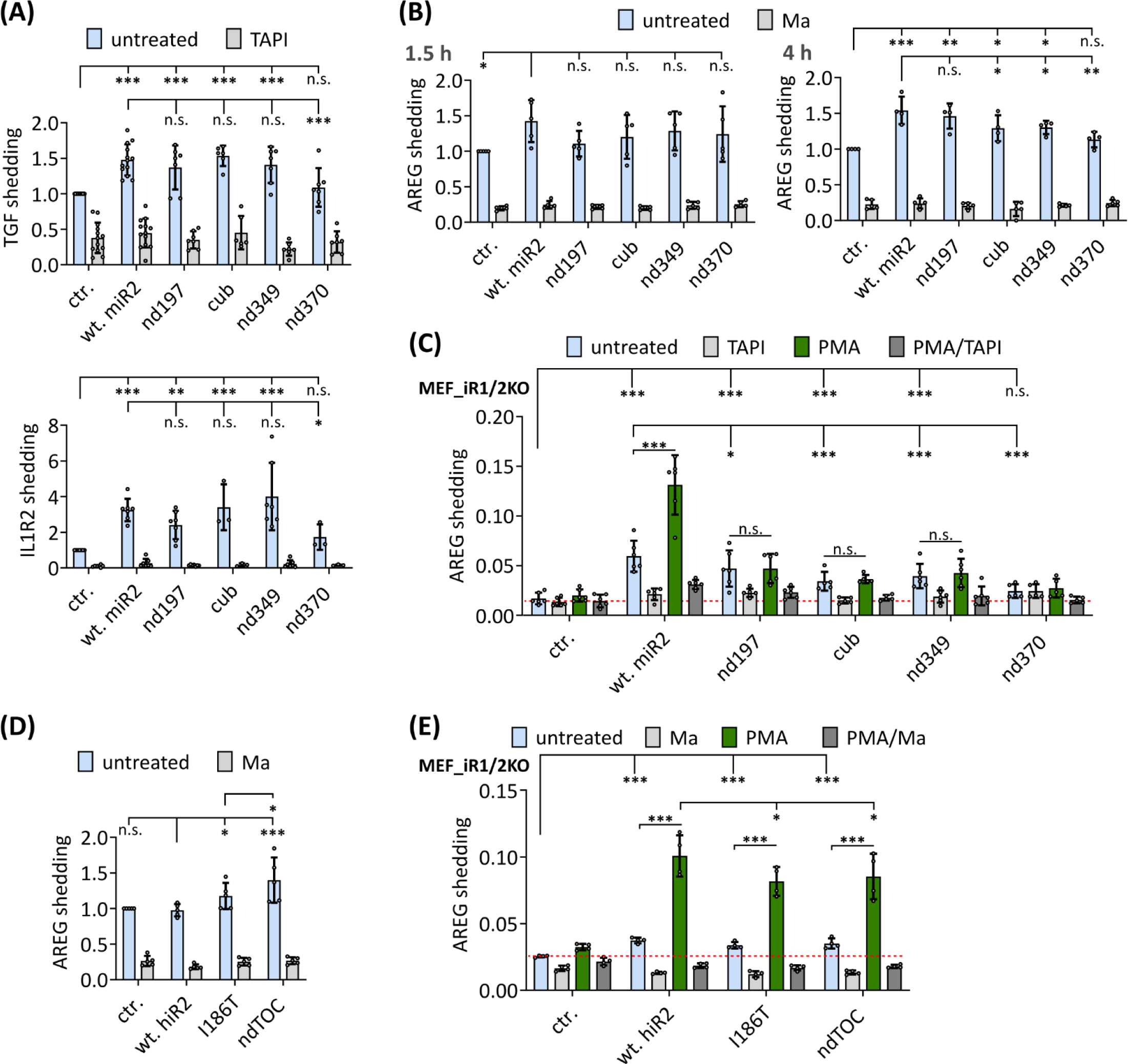
N-terminal mutations in iRhom2 dysregulate ADAM17-mediated shedding. **(A-E)**Constitutive ADAM17-mediated shedding activity was assessed by performing an alkaline phosphatase (AP) assay in HEK293 cells (A,B,D) and MEFs deficient for iRhom1 and iRhom2 (C,E) expressing the indicated iRhom2 constructs. The ADAM17 substrate TGFα, IL1R2 or AREG tagged with AP were used. The metalloprotease inhibitors marimastat (10 µM) or the more ADAM17 specific inhibitor TAPI1 (10 µM) were used as negative control. Where indicated, ADAM17 activity was additionally stimulated by the phorbol ester PMA (100 nM). n > 4.

To further examine the effects of large N-terminal deletions in iRhom2 on ADAM17-mediated shedding, we used iRhom1- and iRhom2-deficient mouse embryonic fibroblasts (MEFs)(Christova et al., 2013). These cells lack production of mature ADAM17, resulting in an absence of ADAM17-mediated shedding. However, reintroduction of wt-iRhom2 expression effectively restored ADAM17-mediated shedding (Fig. 4C). Surprisingly, constitutive ADAM17-mediated shedding was also restored for the N-terminal deletions (nd197, cub and nd349), despite barely detectable amounts of mature ADAM17 present (Fig. S3D). However, the shedding activity restored by the N-terminal deletions was significantly lower compared to cells expressing wt-iRhom2 (Fig. 4C). Interestingly, the largest deletion (nd370), which lacked the third phosphorylation site, showed no restored shedding activity, similar to control cells (Fig. 2G; Fig. 4C). Furthermore, we assessed ADAM17-mediated shedding stimulated by the phorbol ester PMA (phorbol 12-myristate 13-acetate). While cells expressing wt-iRhom2 showed a significant increase in shedding, this effect was not observed with iRhom2 deletion mutants (Fig. 4C). This aligns with previous reports indicating that stimulated shedding is primarily driven by phosphorylation at the described phosphorylation sites 1 and 2 (Cavadas et al., 2017, Grieve et al., 2017), which are absent in all deletions (Fig. 2G).

In HEK293 cells, we further investigated the effects of the TOC mutations on AREG shedding. Interestingly, additional expression of human wt-iRhom2 did not significantly increase constitutive AREG shedding (Fig. 4D). This is consistent with our previous findings that overexpression of human iRhom2 promotes shedding to a lesser extent than that of its mouse counterpart (Kahveci-Türköz et al., 2023). Remarkably, iRhom2 has 96% sequence similarity (92% identity) in human and mouse, except for a large insertion at the beginning of the N-terminus of human iRhom2 isoform 1, which was used in this study (Fig. S3A). The discrepancy in shedding efficiency may be attributed to this dissimilarity. Importantly, expression of the TOC mutations significantly increased shedding activity (Fig. 4D), consistent with previous observations (Blaydon et al., 2012, Maney et al., 2015). Strikingly, we observed even higher shedding efficiency with the complete TOC site deletion ndTOC (Fig. 4D), again showing a more severe molecular phenotype than the single point mutations. Moreover, as expected, iRhom2 with TOC site mutations still promote PMA-stimulated ADAM17 shedding activity in iRhom1- and iRhom2-deficient MEFs (Fig. 4E). However, the stimulated shedding activity of the TOC site mutants is slightly but significantly reduced compared to wt-iRhom2, which is in line with reduced levels of mature ADAM17 (Fig. 3C,E; Fig. 4E).

In summary, our results show that the N-terminus differentially regulates constitutive and stimulated shedding and that the cub deletion loses the ability to promote stimulated ADAM17 activity. Importantly, both larger N-terminal deletions and TOC mutations appear to lead to a reduction in mature ADAM17 levels with concurrent higher constitutive ADAM17 activity.

### N-terminal deletions of iRhom2 accelerate turn-over of mature ADAM17

To gain better insight into the molecular phenotypes of N-terminal iRhom2 mutations, we sought to understand the underlying mechanisms. The observed decrease in mature ADAM17 levels can be attributed to either an impaired maturation process or accelerated turnover leading to increased internalisation of ADAM17 and its lysosomal degradation (Fig. 5A). Previous studies have shown that destabilisation of the iRhom-ADAM17 complex by FRMD8 loss leads to increased lysosomal degradation (Künzel et al., 2018, Oikonomidi et al., 2018). However, the effect we observed is FRMD8 independent and has not been studied before. To investigate whether the FRMD8-independent effect of nd197 is due to accelerated degradation, we used the lysosomal inhibitors NH_4_Cl and bafilomycin. The use of these inhibitors for 8 h and 16 h rescued the levels of mature ADAM17 (Fig. 5B,C). After 16 h, the levels of mature ADAM17 in HEK293 cells expressing the cub or nd197 deletion reached those of cells expressing wt-iRhom2 (Fig. 5C), indicating a similar maturation efficiency. Therefore, accelerated turnover is likely the cause of the low mature ADAM17 levels when the deletions are expressed. This is consistent with the reduced total levels of ADAM17 in cells expressing the deletions, as maturation is still occurring, which depletes the pool of immature ADAM17, but the mature ADAM17 is not stable due to accelerated lysosomal degradation.

**Figure 5:**
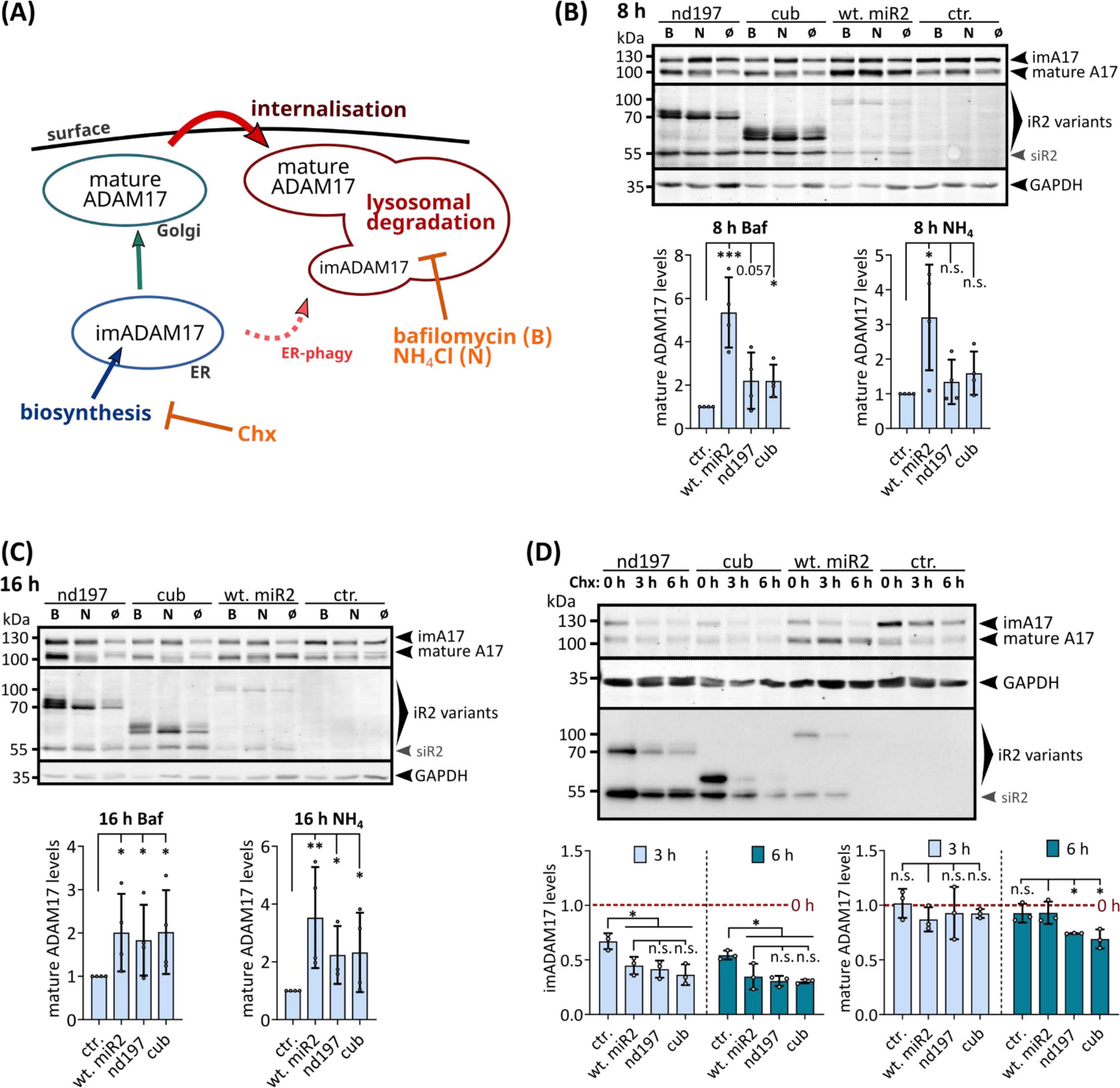
The N-terminal tail of iRhom2 regulates protein turn-over kinetics. **(A)**After biosynthesis in the ER, ADAM17 is transported to the Golgi for maturation with the help of iRhoms. The complex of mature ADAM17 and iRhom is transported to the cell surface. After internalisation, the complex is degraded in the lysosomes. Like all ER proteins, to a certain extent immature ADAM17 can be transported directly to the lysosome by ER-phagy. **(B-C)** To analyse the fate of mature ADAM17 in cells expressing the nd197 and cub deletion, cells were treated with either NH_4_Cl (“N”) or bafilomycin (“B”) for 8 h (B) or 16 h (C) to inhibit lysosomal degradation. Levels of mature ADAM17 were assessed by densitometric measurements and calculation of the ratio between mature ADAM17 and the loading control GAPDH, which is independent of lysosomal degradation. (n = 4) **(D)** To assess the turn-over rate of ADAM17 a cycloheximide-based (Chx) pulse-chase experiment was performed. Reduction of ADAM17 levels was assessed by densitometric measurements and normalisation to 0 h Chx treatment (set to 1). (n = 3).

To further analyse the impact of the iRhom2 N-terminus on the turnover kinetic of ADAM17, we performed a cycloheximide-based pulse-chase experiment (Fig. 5A). We observed a rapid decrease in immature ADAM17 levels after inhibition of biosynthesis, as the maturation process continued and subsequently immature ADAM17 levels declined until presumably no iRhom is left in the ER (Fig. 5D). The presence of wt-iRhom2 facilitated this effect, significantly enhancing the reduction due to increased ADAM17 transport to the Golgi and subsequent maturation. We found that nd197 and cub deletions had the same effect (Fig. 5D), indicating that maturation process remained unaffected by the N-terminal deletions.

Furthermore, the levels of mature ADAM17 decreased more rapidly in cells expressing the deletion than in control cells or cells expressing wt-iRhom2 (Fig. 5D) indicating accelerated degradation of mature ADAM17 in the presence of N-terminal deletions. For comparison, we next studied the influence of TOC mutations on ADAM17 stability. 8 h of bafilomycin-mediated lysosomal inhibition rescued the levels of mature ADAM17 in cells expressing the TOC mutant to the same level as cells with wt-human iRhoms (Figure S3E). This suggests that the same amount of mature ADAM17 is generated in cells expressing TOC mutants or wt-human iRhom2. Lysosomal degradation of mature ADAM17 is slightly higher in cells with TOC mutations than in cells expressing wt-iRhom2, resulting in slightly lower levels of total mature ADAM17 and less mature ADAM17 on the cell surface, as described above (Figure 3C,E). However, this slightly accelerated degradation does not seem to be strong enough to detect significant differences with the cycloheximide-based pulse-chase experiments (Figure S3F).

Taken together, these results demonstrate the regulatory role of the N-terminus in controlling the stability and turnover kinetics of mature ADAM17, but not immature ADAM17. In addition, the effects of larger deletions are more severe compared to mutations or deletions at the TOC site, suggesting that the stability of mature ADAM17 is only slightly affected by the TOC site, but is more dependent on the interplay of other sites in the iRhom2 N-terminus. As there is no difference between nd197 and cub, this mechanism appears to be FRMD8 independent.

### The iRhom2 N-terminal tail is a key determinant for regulating ADAM17 binding

The interaction between iRhom and ADAM17 is a crucial requirement for their stability. Although the transmembrane regions and extracellular domains are known determinants of the binding between iRhom and ADAM17 (Cavadas et al., 2017, Grieve et al., 2017, Li et al., 2017, Düsterhöft et al., 2021, Kahveci-Türköz et al., 2023), we nevertheless wondered whether the N-terminus could play a role in this interaction. By co-immunoprecipitation, we analysed the binding between ADAM17 and iRhom2. In both HEK293 cells and iRhom1- and iRhom2-deficient MEFs, we observed a significant decrease in binding between ADAM17 and all deletions tested (nd197, cub, nd349 and nd370) compared to wt-iRhom2 (Fig. 6A,B). However, the overall decrease in cellular ADAM17 levels could distort the results. To address this issue, we engineered iRhom2 variants fused to the ER retention sequence KDEL, which inhibits ADAM17 transport, maturation, and subsequent accelerated turnover of the iRhom2-ADAM17 complex (Fig. S4A) (Düsterhöft et al., 2021). In fact, even in this setting, we found a significant reduction in ADAM17 binding for iRhom2 deletions compared to wt-iRhom2 (Fig. 6C; Fig. S4B). Interestingly, we detected an additional significant reduction in ADAM17 binding for deletions larger than nd197, indicating multiple regions in the N-terminus that regulate ADAM17 binding (Fig. 6A,C). This observation is consistent with co-immunoprecipitation experiments using lysosomal inhibitors, where inhibition of lysosomal degradation significantly increased the observable intact iRhom2-ADAM17 complex with nd197, but not with cub deletion (Fig. S4C). Under lysosomal inhibition, the precipitated complexes with nd197 and cub showed increased amounts of mature ADAM17 indicating more rapid degradation when in complex with these deletions (Fig. 6D,E). Importantly, we did not observe a reduction in binding efficiency of the TOC site mutants (Fig. 6F).

**Figure 6:**
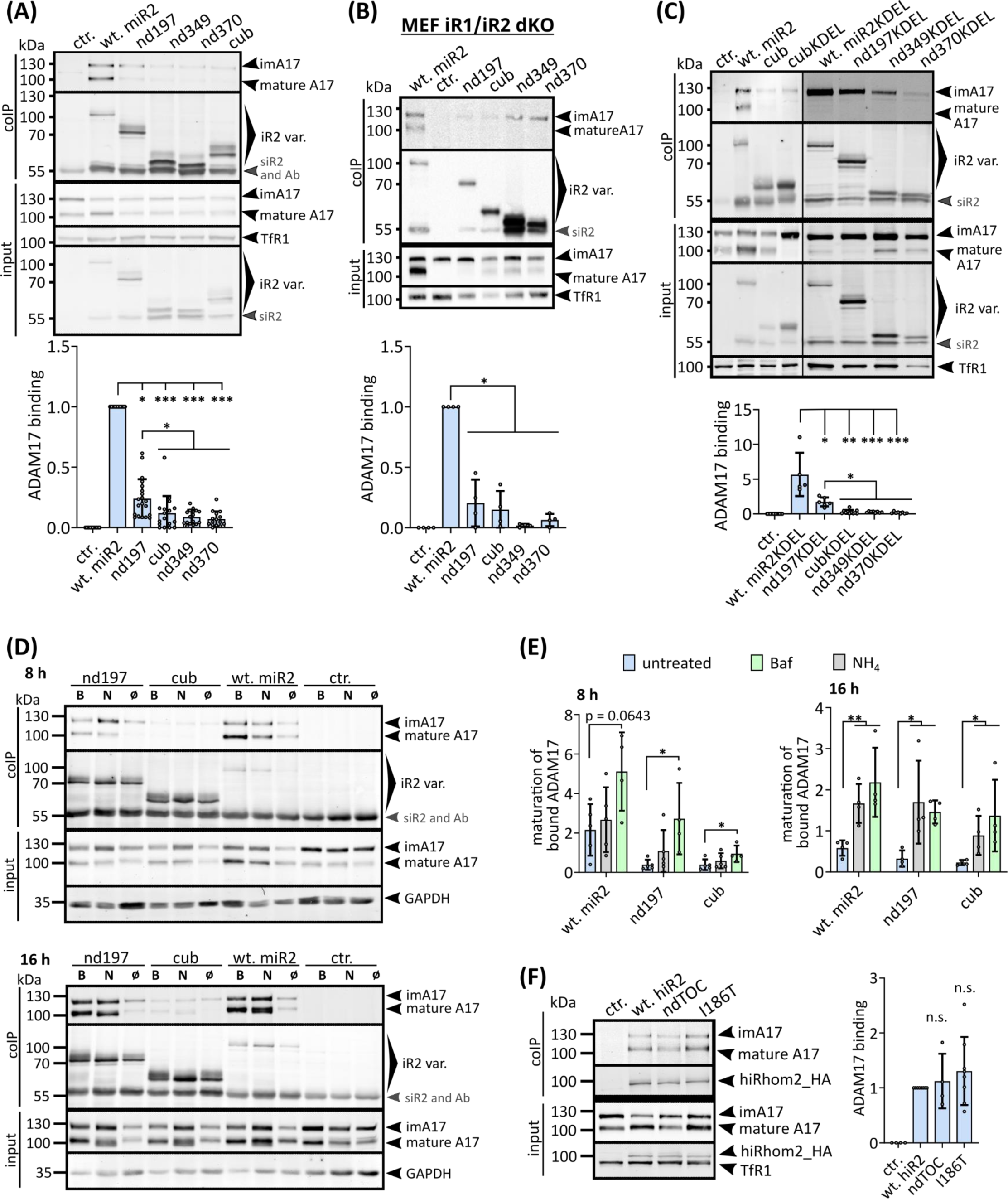
The N-terminus of iRhom2 regulates ADAM17 interaction. To analyse interaction between ADAM17 and the indicated iRhom2 variants, coIPs were performed using the iRhom variants (with HA tag) as bait. Immunoblotting and subsequent densitometric measurements were used to quantitatively analyse iRhom2-ADAM17 binding (ratio of co-precipitated ADAM17 and precipitated iRhom2) and normalised to wt-iRhom2. **(A)** HEK293 stably expressing indicated iRhom2 variants or GFP (ctr.) were used (n > 17). **(B)** MEFs from mice deficient for iRhom1 and iRhom2 stably expressing indicated iRhom2 variants or GFP (ctr.) were used (n = 4). **(C)** HEK293 expressing indicated iRhom2 variants or GFP (ctr.) fused to the ER-retention motif KDEL were used (n > 5). **(D) – (E)** To analyse the fate of mature ADAM17 specifically in the iRhom2-matureADAM17 complex containing either the nd197 deletion, cub deletion or wt-iRhom2, cells were treated with either 40 mM NH4Cl (N) or 500 nM bafilomycin (B) for 8 h or 16 h to inhibit lysosomal degradation. Maturation levels of bound ADAM17 was assessed by densitometric measurements and calculation of the ratio between mature ADAM17 and immature ADAM17. The quantitative analysis of ADAM17 binding can be found in Fig. S4C. (n = 4). **(F)** HEK293 stably expressing indicated human iRhom2 variants or GFP (ctr.) were used for coIP experiments as describe above. (n > 4).

In summary, our results clearly demonstrate the crucial role of the N-terminus for the binding affinity between iRhom2 and ADAM17, which in turn influences the cohesion and stability of the iRhom2-ADAM17 complex. Remarkably, there appear to be two parallel mechanisms regulating the ADAM17 interaction via the N-terminus: a clearly FRMD8-independent mechanism and a potential FRMD8-dependent mechanism. Importantly, this ADAM17 interaction function is lost in the larger cub deletion but not when only the TOC site is disrupted (Fig. 7).

**Figure 7:**
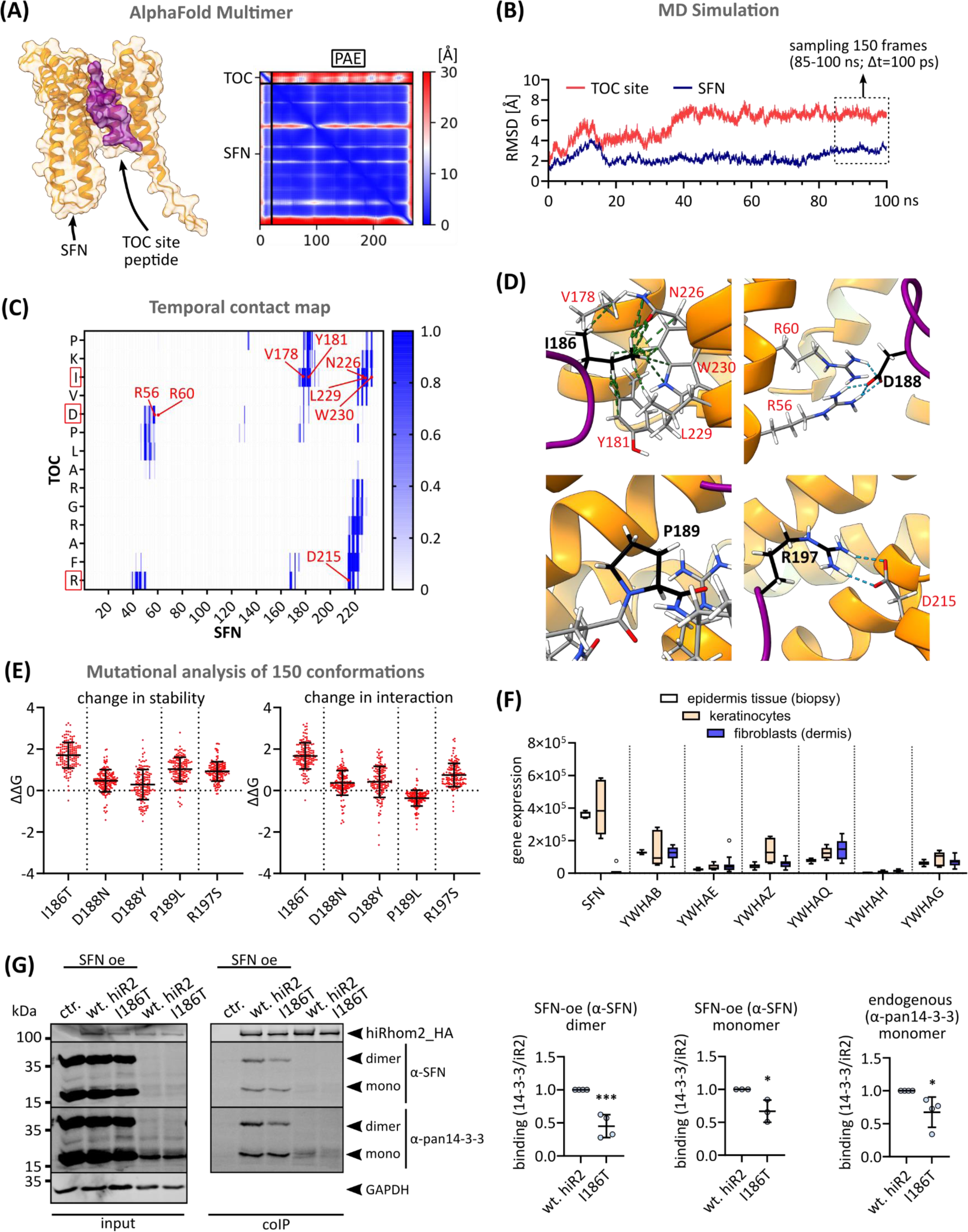
The TOC site binds 14-3-3 proteins in a phosphorylation-independent manner. **(A)**The complex consisting of the TOC site peptide (P184 to R197) and 14-3-3σ/stratifin (SFN) was modelled with AlphaFold Multimer. The highest ranked model and its PAE score are shown. **(B)** The highest ranked AlphaFold model of the SFN-TOC site complex was used for molecular dynamics simulation. RMSD changes during the simulation run are shown. 150 frames/conformations between 85 ns and 100 ns (Δt = 100 ps) were extracted for subsequent analysis. **(C)** Combined Cα contact map of all 150 sampled conformations indicates the frequency of contacts between SFN and TOC site over time. Identified interactions between positions of TOC mutants and residues in SFN are labelled. **(D)** Visualisation of interactions between pathologically relevant residues in the wt TOC site with residues of SFN: cartoon representation of conformation at 92 ns of MD simulation. I186 interacts with a hydrophobic cleft in SFN. D188 and R197 form salt bridges. P189 shows no interaction. **(E)** Changes in overall complex stability and binding energies were calculated for the pathological TOC mutations for all 150 conformations: ΔΔG<0 indicate improved stability/binding; ΔΔG>0 indicate decreased stability/binding. **(F)** Gene expression levels of 14-3-3 proteins in epidermis and primary keratinocytes and primary fibroblasts. **(G)** To analyse the interaction between 14-3-3 proteins and the indicated iRhom2 variants, coIPs were performed using the indicated iRhom variants (with HA tag) as bait. SFN was additionally overexpressed (oe) in the indicated samples. Immunoblotting and subsequent densitometric measurements were used to quantitatively analyse iRhom2-14-3-3 protein binding (ratio of co-precipitated 14-3-3 protein and precipitated iRhom2) and normalised to wt iRhom2. Notably, the dimeric form of oe SFN appears to be somewhat resistant to SDS.

### The TOC site represents a phosphorylation independent 14-3-3 binding site

Using AlphaFold Multimer complex modelling, we screened the TOC site for interaction with putative iRhom2 binding partners that we and others had previously reported (Cavadas et al., 2017, Grieve et al., 2017, Düsterhöft et al., 2021). Thereby we identified 14-3-3σ/stratifin (SFN) as a potential interactor for the TOC site (Fig. 7A). To analyse the dynamic nature of the TOC-site-SFN complex, we performed molecular dynamics simulations and sampled 150 conformations within the RMSD (root-mean-square deviation) equilibrium (Fig. 7B). A contact map based on this conformational assembly revealed the interacting residues between the TOC site and SFN (Fig. 7C,D). Specifically, I186 interacts with a hydrophobic cleft in SFN, while D188 and R197 form salt bridges with SFN residues (Fig. 7D). P189 shows no direct interaction but appears to maintain a local structure important for overall binding with SFN (Fig. 7D,E). Strikingly, D188 forms a salt bridge with R56 and R60 of SFN, which are part of the region that normally facilitates interaction with phosphorylated serine residues of 14-3-3 clients. Thus, D188 appears to mimic a phosphoserine residue, facilitating a phosphorylation-independent mode of 14-3-3 binding. The consequences of the pathological TOC mutations on the complex stability and interaction energies were assessed and, as expected, showed a profound effect on the complex integrity (Fig. 7E). Based on these findings, we investigated whether overexpressed SFN, which is predominantly expressed in skin and undetectable at endogenous levels in HEK293 cells (Fig. 7F,G), shows altered interaction efficiency between wt iRhom2 and the I186T TOC mutant. Indeed, using coIPs, we found that the I186T TOC mutant showed reduced interaction with SFN compared to wt iRhom2 (Fig. 7G). These data are in line with our *in silico* analysis and confirm the role of I186 within TOC for efficient SFN binding. Using a pan-14-3-3 antibody, we also found that other 14-3-3 isoforms expressed endogenously interact with the TOC site (Fig. 7G).

Overall, the TOC site serves as a phosphorylation-independent 14-3-3 binding site and this interaction is disrupted by pathological TOC mutations.

## Discussion

The iRhom2-ADAM17 sheddase complex is vital for both inflammatory signals (including TNFα release) and EGFR signalling (via growth factor release such as AREG and TGFα). iRhom2 is a key regulator for ADAM17’s trafficking, maturation, activity, and stability. Understanding its regulation is crucial for the molecular biology of ADAM17-associated diseases and, consequently, for potential therapies targeting these pathologies. These include chronic inflammation and cancer pathologies, such as the autosomal dominant hereditary disorder Tylosis Oesophageal Carcinoma (TOC) (Blaydon et al., 2012, Düsterhöft et al., 2019). TOC is caused by mutations in the cytosolic N-terminus of iRhom2, resulting in increased ADAM17 activity, elevated growth factor release and increased EGFR signalling. Mice with the cub (curly-bare) deletion exhibit a similar phenotype to mice with TOC mutations, and both phenotypes can be rescued by knocking out amphiregulin (Hosur et al., 2017, Hosur et al., 2018b). This suggests a shared molecular mechanism between the large cub deletion and TOC single point mutations. However, studies on the cub deletion have yielded conflicting findings regarding whether it has a positive or negative effect on ADAM17 shedding activity (Hosur et al., 2014, Siggs et al., 2014, Maney et al., 2015, Grieve et al., 2017). In contrast, TOC mutations in iRhom2 clearly increase ADAM17-mediated shedding (Blaydon et al., 2012, Brooke et al., 2014), which is consistent with our results. In addition to TOC single point mutations, we used a complete deletion of the TOC site, resulting in an even more pronounced effect.

We identified the TOC site as a phosphorylation-independent 14-3-3 protein binding site that appears to have a restrictive effect on constitutive ADAM17 activity, which is disrupted by the pathological TOC mutations (Fig. 8). 14-3-3 proteins, as phosphoserine and phosphothreonine binding proteins, regulate most cellular processes. With widespread expression in tissues, they serve as universal scaffolds, influencing the structure and function of diverse protein targets in key cellular signalling pathways (Obsilova and Obsil, 2022). Although the majority of reported 14-3-3 binding motifs involve phosphorylation, there are a few non-canonical examples. In particular, the L**D**LA box in the exoenzyme S of *Pseudomonas aeruginosa* and the WL**D**LE motif in the artificial peptide R18 show high affinity to 14-3-3 proteins (Obsilova and Obsil, 2022). These motifs consistently feature an aspartate residue surrounded by hydrophobic residues, which bears a striking resemblance to the IV**D**PL motif found within the TOC site. Our finding is also consistent with previous results that the TOC site may also facilitate binding to keratin (Maruthappu et al., 2017). 14-3-3 proteins are known to interact with keratins (Ku et al., 2002) and may therefore act as an adapter between iRhom and keratins.

**Figure 8:**
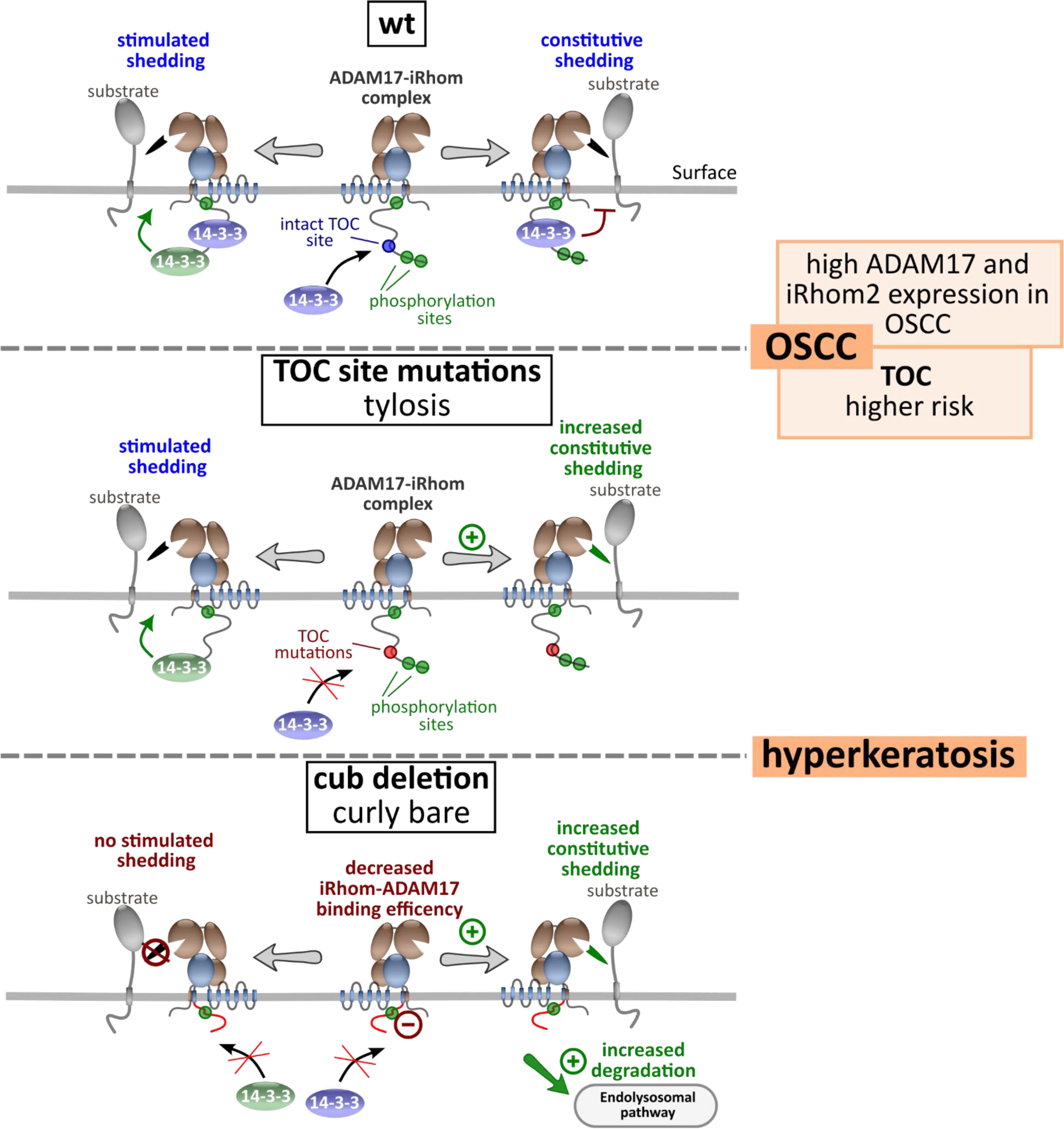
The iRhom2 N-terminal tail regulates ADAM17 binding and shedding activity. Wt iRhom2 plays a critical role in regulating the transport, maturation, constitutive and simulated shedding activity of ADAM17, ensuring proper control of downstream signalling pathways such as EGFR activation. Conversely, disruption of the TOC site in the N-terminal tail of iRhom2 results in loss of phosphorylation-independent 14-3-3 binding and appears to result in dysregulated higher constitutive shedding activity. Hyperkeratosis and increased risk of OSCC, which also shows higher iRhom2 and ADAM17 expression, are known consequences of TOC mutations. In the case of the cub deletion, not only is the TOC site disrupted, but efficient binding to ADAM17 is also impaired, accelerating ADAM17 turnover and lysosomal degradation. In addition, the cub deletion loses its ability to promote stimulated ADAM17 shedding activity. It is likely that the lack of phosphorylation-dependent 14-3-3 binding plays a key role in this process.

In contrast, the larger cub deletion, which includes the entire TOC site, has more molecular consequences (Fig. 8): 1) The loss of stimulated ADAM17 activity is consistent with the loss of N-terminal phosphorylation sites. 2) Large deletions of the N-terminus reduce the efficiency of iRhom-ADAM17 interaction and accelerate ADAM17 degradation, resulting in reduced levels of mature ADAM17 and reduced ADAM17 surface localisation.

Previously identified phosphorylation-dependent 14-3-3 binding to the iRhom N-terminus appears necessary for stimulated shedding (Cavadas et al., 2017, Grieve et al., 2017)(Fig. 8). In contrast, constitutive shedding may be attenuated by efficient 14-3-3 binding to the TOC site, which would be consistent with previously reported ADAM17 hyperactivation (Blaydon et al., 2012, Maney et al., 2015, Hosur et al., 2017). Both the cub deletion and the TOC single point mutations still show substantial amounts of constitutive ADAM17 shedding activity. It is important to note that while the cub deletion results in less ADAM17 activity being measured compared to wt iRhom2, it also results in a profound reduction of mature ADAM17 on the cell surface. Taking this into account, the shedding activity per ADAM17 molecule appears to be up-regulated, while the overall shedding activity at the cellular level appears to be reduced.

Interestingly, the largest deletion we used, nd370, which is also missing a third key phosphorylation site and putative 14-3-3 binding site (Fig. 2G)(Cavadas et al., 2017, Grieve et al., 2017), additionally shows reduced constitutive shedding. Taken together, these suggest that there are distinct signalling pathways which differentially regulate 14-3-3 binding and, in turn, constitutive and stimulated ADAM17 activity. More studies are needed to dissect these complex mechanisms.

Notably, recent studies have also highlighted the critical role of the KRAS-ERK1/2 pathway in the shedding activity of the iRhom2-ADAM17 complex (Cavadas et al., 2017, Grieve et al., 2017, Sieber et al., 2022). KRAS-G12 mutants in tumour cells harbouring iRhom2 with TOC mutations further enhance constitutive ADAM17-mediated shedding (Sieber et al., 2022). Interestingly, an ERK2 binding site was recently identified within the N-terminus upstream of phosphorylation sites 1 and 2 (Fig. 2G)(Schumacher et al., 2023). Since this ERK2 interaction site is not present in cub, it appears to be more relevant for stimulated shedding rather than constitutive shedding.

The N-terminus of iRhoms emerges as a critical determinant for the stability and activity of the iRhom-ADAM17 complex. This is again supported by our data and was recently also confirmed *in vivo*: mice with the cub mutation show decreased ADAM17 levels in different tissues (Rabinowitsch et al., 2023). Recent findings highlight the significance of FRMD8-binding in regulating iRhom stability and, consequently, ADAM17 stability and functions (Künzel et al., 2018, Oikonomidi et al., 2018). While the loss of FRMD8 binding may contribute to the molecular consequences observed in cub deletions, our observations demonstrate similar effects on mature ADAM17 stability in the nd197 deletion, which retains the ability to bind FRMD8. This suggests that the N-terminus exerts a stabilising effect on the iRhom-ADAM17 complex, largely independent of FRMD8: When these parts of the iRhom2 N-terminus, which are not involved in FRMD8 binding, are deleted, accelerated turnover still occurs. Notably, TOC mutants also show an effect on the stability of mature ADAM17, but to a lesser extent.

Our study reveals that not only the transmembrane helix 1 and the IRHD, as previously described (Li et al., 2017, Tang et al., 2020, Düsterhöft et al., 2021, Kahveci-Türköz et al., 2023), but also the iRhom2 N-terminus, contributes to ADAM17 binding. Through our ER-trapping approach, we have demonstrated that the reduced binding of ADAM17 is not attributable to decreased ADAM17 levels. Notably, the anterior half of the N-terminus (nd197) emerges as particularly crucial for efficient ADAM17 interaction. However, larger deletions like the cub deletion (Δ268) result in an additional significant reduction in binding efficiency, potentially mediated by FRMD8.

We postulate that while the binding affinity to ADAM17 is significantly reduced, iRhom2 N-terminal deletions such as cub retain the ability to bind immature ADAM17 in the ER, facilitating efficient transport through the secretory pathway and normal maturation of ADAM17. During maturation, the proteolytic removal of the ADAM17 prodomain may further decrease binding affinity, as our recent study identified the prodomain as an additional binding determinant between the ADAM17 ectodomain and the IRHD (Kahveci-Türköz et al., 2023). The reduced cohesion of the iRhom2-matureADAM17 complex on the cell surface appears to induce instability and accelerated turnover, culminating in lysosomal degradation (Fig. 8).

In our study, we focused on the molecular functions of the iRhom2 N-terminus to unravel the intricate regulatory system of the iRhom2-ADAM17 complex. It is important to note that our study did not explore the integration of these mechanisms into a more complex *in vivo* system. Importantly, ADAM17 activity at the cellular level *in vivo* is dependent on cell type-specific endogenous protein levels of iRhom1, iRhom2 and ADAM17, which will also influence the manifestations of pathological changes in the iRhom-ADAM17 axis (Rabinowitsch et al., 2023). Here it is also important to consider that ADAM17 substrate specificity appears to be differentially regulated by iRhom1 and iRhom2 (Maretzky et al., 2013, Tang et al., 2020). In addition, iRhom functions independent of the ADAM17 pathway may also be involved in the manifestation of TOC and cub pathologies. For example, the N-terminus of iRhom can be cleaved and translocated into the nucleus where it exerts incompletely understood effects on gene expression (Dulloo et al., 2022, Zanotti et al., 2022). It is also critical to recognise that altered ADAM17 activity can also initiate feedback loops that further modulate its own activity, including the induction of ADAM17 activity by activation of the EGF receptor pathway (Grieve et al., 2017, Kubo et al., 2022, Sieber et al., 2022) and induction of gene expression of ADAM17, iRhoms and other regulators through various pathways including TNFα signalling (Arcidiacono et al., 2018, Babendreyer et al., 2020, Giese et al., 2021, Kubo et al., 2022). In addition, exogenous stimuli and cell stimulation, for example by LPS, also affect ADAM17 activity and gene expression of iRhom2, ADAM17 and its substrates (Arndt et al., 2011, McIlwain et al., 2012, Lu et al., 2017, Düsterhöft et al., 2021).

As the TOC mutations affect iRhom2, it is important to note that high iRhom2 expression is often predominantly associated with immune cells and inflammatory conditions. This role of iRhom2 becomes apparent when considering that mice lacking iRhom2 have a seemingly healthy phenotype but show impaired immune responses (Adrain et al., 2012, Christova et al., 2013, Issuree et al., 2013). However, the high level of iRhom2 gene expression we observed in keratinocytes highlights the important role of iRhom2 in skin homeostasis. This would be consistent with the TOC skin hyperkeratosis phenotype and, in contrast, the reduced thickness of stress calluses in iRhom2 KO mice described previously (Fig. 8)(Blaydon et al., 2012, Brooke et al., 2014, Maney et al., 2015, Maruthappu et al., 2017). Notably, in patients with TOC mutations, tylosis manifests between the ages of 7 and puberty, while cancer develops much later in life (Ellis et al., 2015), hinting at a cumulative effect over time. This is in line with our finding that disruption of the TOC site leads to a gradual effect on iRhom2-ADAM17 complex functions: The larger deletion of the TOC site has a stronger effect on ADAM17 activity than single point mutations.

Furthermore, we found a significant increase in iRhom2 and ADAM17 gene expression in oesophageal squamous cell carcinoma (OSCC) compared to healthy oesophageal squamous tissue. This indicates that the iRhom2-ADAM17 complex is a potential tumour marker for OSCC, which seems to be in line with the fact that patients with TOC mutations, which increases ADAM17 activity, have a higher risk of developing OSCC (Fig. 8).

Overall, our results shed new light on the regulation of the iRhom2-ADAM17 complex by the N-terminal tail of iRhom2. The intrinsically disordered nature of the N-terminus makes it an extensive hub for protein binding and regulation. The molecular functions highlighted in our study warrant further analysis and could potentially serve as viable targets for modulating ADAM17 activity. They could offer promising avenues for combating ADAM17-associated pathologies such as cancer or chronic inflammation.

## Methods

### Bioinformatics

Multiple sequence alignments were generated by utilising Clustal Omega (Sievers et al., 2011). Secondary structure prediction and analysis of intrinsically disordered regions were done with IUPred3 (Meszaros et al., 2009, Erdos et al., 2021), NetSurfP-2.0 (Klausen et al., 2019) and ANCHOR2 (Dosztanyi et al., 2009, Meszaros et al., 2009, Meszaros et al., 2018). The following sequences were used (UniProtKB entries): Q6PJF5 (human iRhom2) and Q80WQ6 (murine iRhom2). Nuclear localisation sequence analysis was performed with NLStraddamus (Nguyen Ba et al., 2009).

To identify conserved sequence regions, the tool CONYAR (<u>Co</u>nserved <u>Y</u>ou <u>Ar</u>e) was employed to perform high-throughput comparison of as many sequences of a target protein as possible (Kahveci-Türköz et al., 2023).

### Structural modelling and molecular dynamics simulations

*Ab initio* structure prediction of the cytosolic iRhom2 N-terminus was done with AlphaFold2 and AlphaFold Multimer (Jumper et al., 2021) utilising ColabFold (Mirdita et al., 2022). Structural modelling using the deep learning algorithm AlphaFold2 was done without homology templates. MMseq2 was used as the multiple sequence alignment. Structural modelling was performed in 12 iterations. ChimeraX was used to visualise the structures (Pettersen et al., 2021).

Molecular dynamics simulation was used to analyse the dynamics of the SFN-TOC-site complex over time. The highest ranked AlphaFold model of the complex was used. The system was immersed in TIP3P water in a cubic box with 1.0 nm padding and neutralised with 150 NaCl. The MD simulation was performed using GROMACS2020.6 (Pronk et al., 2013) with the AMBER99SB-ILDN force field (Lindorff-Larsen et al., 2010). The system was first energy minimised. Equilibration MD simulations were performed with periodic boundary conditions. The temperature was equilibrated using an NVT ensemble for 1 ns followed by an NPT ensemble for 1 ns to equilibrate the pressure using the v-rescale temperature and Parrinello−Rahman pressure coupling method during production. Finally, the production MD simulations were performed for 100 ns (300 K and 1 atm). Between 85 ns and 100 ns, which was within RMSD equilibrium, 150 conformations/frames were sampled (Δt = 100 ps). Biopython (Cock et al., 2009) was used to calculate the distances between Cα within this conformational compilation of each frame and to generate a temporal contact map (cutoff <12 Å) with the frequency of contacts within the sampled 15 ns. FoldX (v5) PSSM and Position Scan (Van Durme et al., 2011) were used to analyse the influence of single point mutations in the sampled conformations on binding and overall stability, respectively.

### In silico gene expression analysis

We accessed Affymetrix Human Genome U133 Plus 2.0 Array platform data from public repositories, curated by the GENEVESTIGATOR platform (Immunai, USA)(Hruz et al., 2008). The GENEVESTIGATOR software was utilised for gene expression meta-analysis of indicated genes in various tissues, biopsies, and primary cells (assessed on 20 May 2023). To analyse gene expression patterns under physiological conditions, only samples from healthy and untreated tissues were included. Similarly, for assessing baseline gene expression in cancerous tissue, only untreated samples were considered.

List of used experiment IDs for indicated tissue and primary cells: a) murine heart: MM-00170 (Muchir et al., 2007), MM-00253 (Dufour et al., 2007), MM-00257 (Zhao et al., 2007), MM-00262 (Alcendor et al., 2007), MM-00300 (Ramchandran et al., 2006), MM-00322, (Thorrez et al., 2008), MM-00330 (Lattin et al., 2008), MM-00368 (Barger et al., 2008), MM-00384 (Wong et al., 2010), MM-00476 (Abd Alla et al., 2016), MM-00485 (Oka et al., 2011), MM-00492 (Wang et al., 2016), MM-00497 (Shuai et al., 2011), MM-00507 (Shuai et al., 2011), MM-00561 (Ricard et al., 2010), MM-00834 (Traister et al., 2013), MM-00841 (Mercader et al., 2012), MM-01538 (Gambino et al., 2013), MM-01642 (Lucas et al., 2014), MM-01875 (Lucas et al., 2014); b) murine primary fibroblast (dermis): MM-00159 (Vallender and Lahn, 2006), MM-00180 (Vallender and Lahn, 2006), MM-00627 (Garinis et al., 2009), MM-00724 (Garinis et al., 2009); c) murine primary keratinocytes: MM-01215 (Allende et al., 2013); d) healthy human epidermis tissue (biopsy): HS-02924 (Esaki et al., 2015); e) human primary keratinocytes (untreated): HS-02625 (Yunoki et al., 2018), HS-02787 (Busch et al., 2008), HS-02490 (Swindell et al., 2012), HS-00220 (Sa et al., 2007); f) human primary keratinocytes (treated with TNFα, IFNα, IFNγ or IL4; 24 h): HS-02490 (Swindell et al., 2012); g) human primary keratinocytes (treated with 10 ng/ml I IFNγ; 96 h): HS-02625 (Yunoki et al., 2018); h) human primary fibroblasts (untreated): HS-00393, HS-00481 (Chen et al., 2008), HS-00805 (Verkerk et al., 2009), HS-00928 (Wang et al., 2012), HS-00963, HS-00989 (Strunnikova et al., 2009), HS-01109 (Visser et al., 2012), HS-01279 (Marchina et al., 2014), HS-01311 (Henrichsen et al., 2011), HS-01326 (Voets et al., 2012), HS-02421 (Smith et al., 2008); i) primary artery smooth muscle cells (heart): HS-00478 (Shalhoub et al., 2010); j) primary bronchial smooth muscle cells: HS-00039 (Bosse et al., 2007); k) healthy oesophagus squamous tissue (biopsy): HS-01298 (Wang et al., 2013), HS-00217, HS-00017 (Roth et al., 2006), HS-01525 (Sato et al., 2013); squamous cell carcinoma (grade G1): HS-01298 (Wang et al., 2013); oesophagus adenocarcinoma: HS-00002; HS-01298 (Wang et al., 2013); Barrett’s oesophagus (pre-malignant): HS-01298 (Wang et al., 2013); primary macrophages: HS-01265; primary granulocytes: HS-00250; primary leukocytes: HS-00261; primary monocytes: HS-01403.

### Cloning

The cloning procedures were performed as previously described (Düsterhöft et al., 2021): NEBuilder HiFi DNA Assembly Master Mix (NEB, E2621L) was utilised to produce different plasmids with the desired inserts according to the manufacturer’s instructions. All iRhom constructs were cloned into the pMOWS backbone (Ketteler et al., 2002, Düsterhöft et al., 2021). Site-directed mutagenesis was performed using overlapping PCR (Ho et al., 1989).

### Cell culture

All cells were cultured in a humidified incubator at 37 °C, 5% CO_2_ in DMEM10% (unless otherwise indicated). DMEM10% consists of DMEM high-glucose (Sigma-Aldrich) supplemented with 10% foetal calf serum (PanBiotech), 100 mg/l streptomycin (Sigma-Aldrich) and 60 mg/l penicillin (Sigma-Aldrich). Production of stable cell lines was done as described before (Düsterhöft et al., 2021). MEFs double deficient for iRhom1 and iRhom2 were obtained from indicated mouse lines as described before (Christova et al., 2013, Düsterhöft et al., 2021). HEK293 cells were purchased from German Collection of Microorganisms and Cell Cultures (GmbH DSMZ-No. ACC 305). Normal human epidermal keratinocytes and human dermal fibroblasts were isolated from human skin samples as described before (Schmitt et al., 2018). Primary keratinocytes were cultivated in Dermalife K – Life Line (Cell Systems LL-0007). Primary fibroblasts were cultivated in DMEM10%.

### Quantitative PCR (RT-qPCR)

mRNA expression levels were measured by quantitative PCR and normalised to the mRNA expression levels of the selected reference genes. Cytochrome c1 (CYC1) and ribosomal protein L13a (RPL13A) were chosen as the most stable reference genes based on the results from the CFX Maestro Software 1.1 (Bio-Rad). RNA was extracted using RNeasy Kit (Qiagen) and quantified photometrically (NanoDrop, Peqlab, Erlangen, Germany). For reverse transcription, the PrimeScriptTM RT Reagent Kit (Takara Bio Europe) was used, according to the manufacturer’s protocol. PCR reactions were performed in duplicates of 10 µl volume comprising 1 µl of cDNA template, 5 µl iTaq Universal SYBR Green Supermix (Bio-Rad), 3 µl H_2_O and 0.5 µl forward and reverse primer. The following primers were used with the primer annealing time given in brackets: CYC1 (forward: AGC TAT CCG TGG TCT CAC C, reverse: CCG CAT GAA CAT CTC CCC ATC; 59 °C), RHBDF1 (forward: GAC AGC CCA CAT CTC TTC AC, reverse: TCC TTG CTC ACT CCA AAC CA; 56 °C), RHBDF2 (forward: CGA TTG ACC TGA TCC ACC, reverse: CAA AGT CTC CGA GCA GTC C; 58 °C) and RPL13A (forward: GCC CTA CGA CAA GAA AAG CG, reverse: TAC TTC CAG CCA ACC TCG TGA; 60 °C). All RT-qPCR reactions were performed on a CFX Connect Real-Time PCR Detection System (Bio-Rad) using the following protocol: 40 cycles of 10 s denaturation at 95 °C, followed by 10 s annealing at the corresponding temperatures and 15 s amplification at 72 °C. PCR efficiency was calculated from the uncorrected RFU values using LinRegPCR version 2020.0 (Ruijter et al., 2009). CFX Maestro Software 1.1 (Bio-Rad) was used for relative quantification.

### Co-immunoprecipitation and enrichment of glycosylated proteins

Precipitation experiments were done as described before (Düsterhöft et al., 2021, Kahveci-Türköz et al., 2023): Cells were lysed in 1 ml lysis buffer (50 mM Tris, 2 mM CaCl_2,_ 137 mM NaCl, 2 mM EDTA, 10 mM 1,10-Phenanthroline, pH 7.5) supplemented with cOmplete™ protease inhibitor (Sigma; 11697498001). The lysates were cleared by centrifugation at 16,000 x g for 20 minutes at 4 °C. For the enrichment of glycosylated proteins, 450 µl of cleared lysate was incubated with 30 µl of Concanavalin A sepharose (Sigma; C9017). For co-immunoprecipitations (coIPs), 450 µl of cleared lysate was incubated with 10 µl of anti-HA magnetic beads (Thermo; 88836). The lysates were incubated with the beads for 90 min. Afterwards, the beads were washed five times with lysis buffer. The beads were heated in 40 µl reducing loading buffer (3% SDS, 16% glycerol, 8% 2-mercaptoethanol, 0.01% bromophenol blue, 0.1 M Tris HCl, pH 6.8) at 65 °C for 20 minutes.

### Western blotting

Samples were subjected to SDS-PAGE and transferred onto PVDF membranes (Millipore, Immobilon-FL). Membranes were blocked with 5% non-fat dry milk in TBS (50 mM Tris, 150 mM NaCl, pH 7.4) for 20 min at room temperature. Primary antibodies were incubated overnight at 4 °C in 0.1% Tween-TBS and 1% BSA. After three washes with 0.1% Tween-TBS, membranes were incubated with secondary antibody for 1 h at room temperature. Additional washing steps included one wash with 0.1% Tween-TBS and three washes with TBS. Protein detection was performed using the Odyssey 9120 imager system (LI-COR) and the ChemiDoc MP Imaging System (Bio-Rad). Band intensities were quantified using Image Studio Lite software (LI-COR). The following primary antibodies were used: αADAM17 (1:1000; Abcam; ab39162), αHA (1:2000; Biolegend; 901502), αTransferrin-receptor (1:2000; Thermo, H68.4, 13-6800), αGAPDH (1:2000; Thermo, GA1R, 15738), αiRhom2 (1:1000; R&D; 996308; MAB10048), α14-3-3sigma (1:5000, Sigma, #PLA0201), α14-3-3pan (1:1000, Cell Signaling Technology, #8312S). The following secondary antibodies were used at the indicated dilutions: DyLight-680-conjugated αmouse (1:20000; Thermo; 35519), DyLight-800-conjugated αrabbit (1:20000; Thermo; 35571), HRP-conjugated goat αmouse and αrabbit (1:20000; Jackson ImmunoResearch Laboratories, Inc).

### Shedding activity - AP-Assay

ADAM17-mediated shedding activity was measured by an alkaline phosphatase (AP)–based assay as described before (Düsterhöft et al., 2021, Kahveci-Türköz et al., 2023). In brief, we used cells transfected with ADAM17 substrates fused to an alkaline phosphatase to analyse shedding activity: The following ADAM17 substrates cloned in pCndA3.1 were utilised: AP-IL1R2, AP-TNFα, AP-AREG. HEK293 cells stably expressing the indicated iRhom2 constructs were transiently transfected with the indicated substrate. In the case of MEFs, the indicated iRhom2 construct combined with the indicated substrate was transiently transfected. Shedding activity was assessed under different conditions: either the metalloprotease inhibitors marimastat (broad-spectrum, 10 µM) (Sigma; M2699) or TAPI1 (ADAM17 preferential inhibitor, 10 µM) (Sigma; SML0739) were used. When indicated, cells were stimulated with 100 nM PMA (Sigma; P1585). DMSO served as the vehicle control. Following treatment, cells were incubated: 90 min (AREG, TGFα), 120 min (IL1R2), or 240 min (AREG) at 37 °C. To evaluate proteolytic activity of ADAM17, we measured alkaline phosphatase (AP) activity in both the supernatant and cell lysates (lysis buffer: 50 mM Tris, 137 mM NaCl, 2 mM EDTA, 10 mM 1,10-phenanthroline, pH 7.5, supplemented with cOmplete™ protease inhibitor). By continuously measuring AP activity at 405 nm using the FLUOstar Optima (BMG LABTECH) after adding a p-nitrophenyl phosphate (PNPP) solution (Thermo; 37620), we calculated the slope representing the change in absorbance at 405 nm per min. ADAM17 activity was determined by calculating the PNPP substrate turnover (AP activity) in the supernatant relative to the total turnover in the supernatant plus cell lysate.

### Flow cytometric analysis

The staining process was conducted using PBS with 0.2% BSA as the assay buffer, and all steps were carried out at 4 °C or on ice. A total of 2×10^5^ cells of interest were incubated with the primary antibody for 1 h. Subsequently, cells were washed twice with 400 μl of assay buffer. The secondary antibody was added, and cells were incubated in the dark for 45 mins. Following two additional washing steps, the fluorescence signal was assessed using flow cytometry (LSRFortessa, BD Biosciences, Heidelberg, Germany), and FlowJo V10 software was employed for data analysis. The geometric mean of fluorescence intensity was determined to assess cell surface localisation. The primary antibodies used, along with their dilutions, were as follows: αADAM17 (1:100; R&D Systems; MAB 9301), αHA (1:500; Biolegend; 901502). The secondary antibody used was Allophycocyanin-conjugated αmouse (1:200; Jackson ImmunoResearch; 115-135-164).

### Lysosomal inhibition

Cell cultures were first grown to approximately 80% confluence. Cells were treated with either bafilomycin A1 (B) or NH_4_Cl (N). An untreated control was also included using DMSO as a vehicle control. The cells were then incubated for 8 h (B: 0.1 µM; N: 20 mM) or 16 h (B: 0.5 μM; N: 40 mM). After incubation, the plates were placed on ice and the medium was discarded. The cells were washed once with PBS and then harvested with 1 ml cold PBS. A 10 µl aliquot was taken from the cell suspension to determine the number of cells for sample adjustment. The harvested cells were centrifuged at 1000 g for 5 min at 4°C and the supernatant was discarded. Cell pellets were used for lysis and coIPs followed by Western blotting as described above.

### Cycloheximide-based pulse chase experiment

Cell cultures were used at approximately 80% confluence. Cells were incubated with cycloheximide (10 μg/ml) for 3 or 6 h. After washing with PBS, the cells were harvested in 1 ml cold PBS and a 10 µl aliquot was taken from the cell suspension to determine the number of cells for sample adjustment. Harvested cells were pelleted and used for lysis followed by Western blotting as described above.

### Statistics

Statistic was done as described before (Düsterhöft et al., 2021): Experiments were repeated at least three times, as indicated in the figure legends. Quantitative data are presented as mean with standard deviation (SD). Statistical analysis was performed using the generalised mixed model analysis (PROC GLIMMIX, SAS 9.4, SAS Institute Inc., Cary, North Carolina, USA), assuming normal, lognormal, or beta distribution, with the day of the experiment considered as a random factor to assess treatment effect differences across the results. Diagnostic tests, including residual analysis and the Shapiro-Wilk test, were utilized. In cases of heteroscedasticity (as determined by the covtest statement), degrees of freedom were adjusted using the Kenward-Roger approximation. All p-values were adjusted for multiple comparisons using the false discovery rate (FDR). Statistical significance was defined as p < 0.05, with the following notation: p: * < 0.05, ** < 0.01, *** < 0.001.

## Declarations

### Ethics approval

This study was conducted according to the principles of the Declaration of Helsinki and approved by the ethics committee of the Medical Faculty RWTH Aachen, Germany (EK 188/14).

## Authors’ contribution

S.D. conceived, designed, and coordinated the study and wrote the manuscript. S.D. and A.L. analysed data and revised the manuscript. S.D., K.B., L.L., S.K.T., C.F., F.S., S.K., D.K., C.G, N.T. and X.G. performed experiments and analysed the results. S.D. conducted protein bioinformatics analysis, structure prediction and MD simulations. A.B. developed the algorithm for high-throughput identification of conserved regions and performed conservation analysis. S.D. and S.K. performed meta-analysis of *in vivo* and *ex vivo* datasets. C.A. generated FRMD8/iTAP knock-out cells. S.H. and J.M.B. isolated primary keratinocytes and fibroblasts. All authors read and approved the final version of the manuscript.

## Abbreviations

a disintegrin and metalloproteinases: (ADAM);
amphiregulin: (AREG);
co-Immunoprecipitation: (coIP);
epidermal growth factor receptor: (EGFR);
FRMD8: (FERM Domain Containing 8)
interferon: (IFN);
interleukin: (IL);
interleukin 1 receptor 2: (IL1R2);
iRhom homology domain: (IRHD);
oesophageal cancer: (OC);
oesophageal adenocarcinoma: (OAC);
oesophageal squamous cell carcinoma: (OSCC);
predicted aligned error: (PAE);
predicted local distance difference test: (pLDDT);
transforming growth factor alpha: (TGFα);
tumour necrosis factor alpha: (TNFα)

## Acknowledgment

We thank Johanna Jakob for cultivation of the primary cells.

---

**Figure S1:**
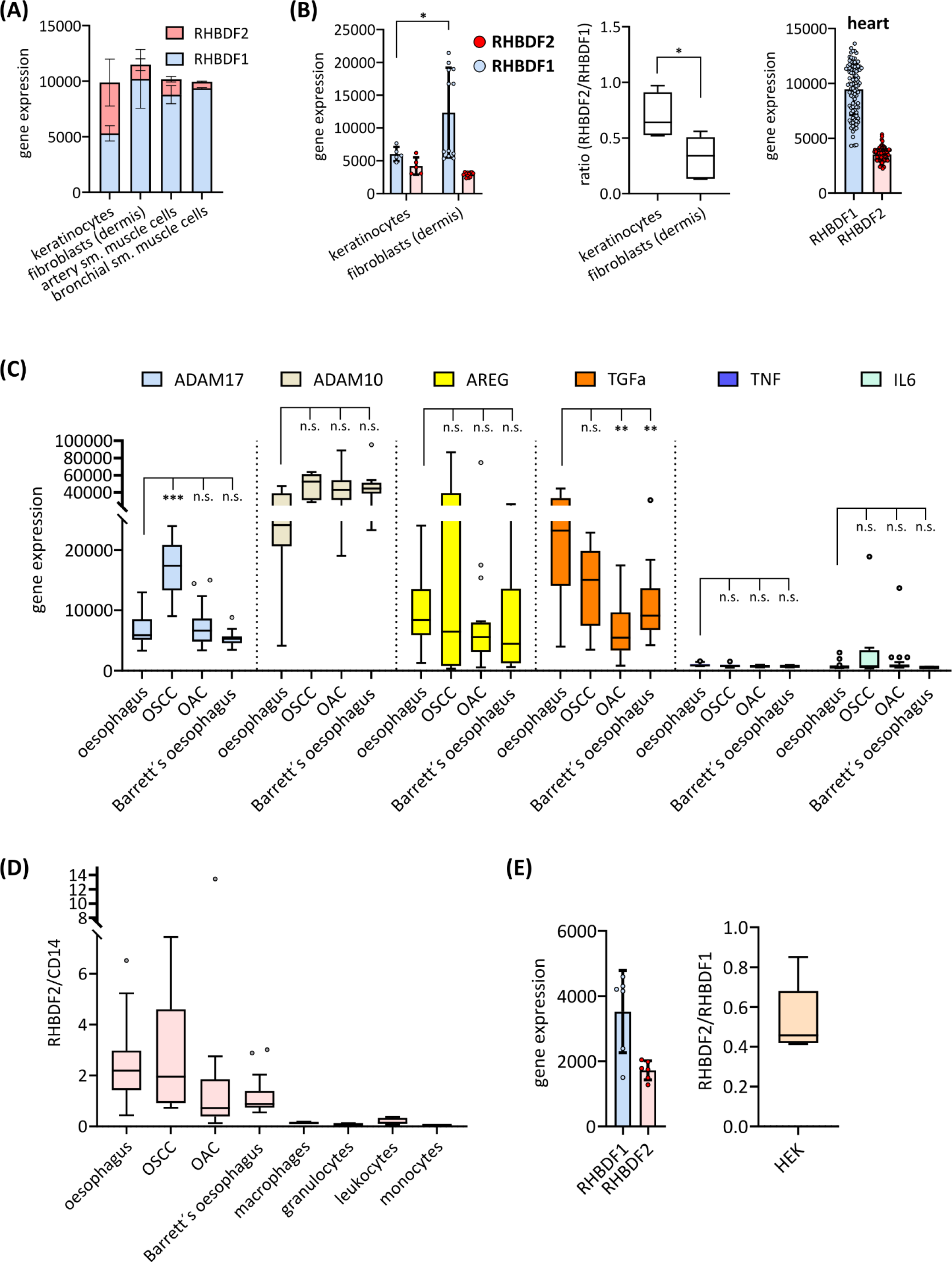
**(A)** Gene expression levels of iRhom1 (RHBDF1) and iRhom2 (RHBDF2) (see Fig. 1C) from primary cell samples are shown as stacked bars, indicating that total iRhom (iRhom1 plus iRhom2) expression may be regulated to a similar overall level in untreated primary cells from healthy tissue. **(B)** Gene expression levels of iRhom1 (RHBDF1) and iRhom2 (RHBDF2) from untreated primary murine keratinocytes from healthy tissue (n = 5), untreated primary murine fibroblasts from healthy dermis tissue (n = 12) and samples from healthy murine heart tissue (n = 98) were analysed. **(C)** Gene expression levels of ADAM17, AREG, TGFα, TNFα and IL6 were analysed in oesophagus and cancerous oesophagus tissues (see Fig. 6C). **(D)** To test whether the increased iRhom2 expression levels were due to the presence of macrophages or monocytes in the samples, the ratio between iRhom2 and CD14 (monocyte and macrophage markers) of the samples was calculated and compared with the values of the indicated immune cell populations (Fig. S2), showing that the measured high iRhom2 levels cannot be explained by the presence of immune cells.**(E)** Gene expression levels of iRhom1 (RHBDF1) and iRhom2 (RHBDF2) fromHEK293 cells (n = 6).

**Figure S2:**
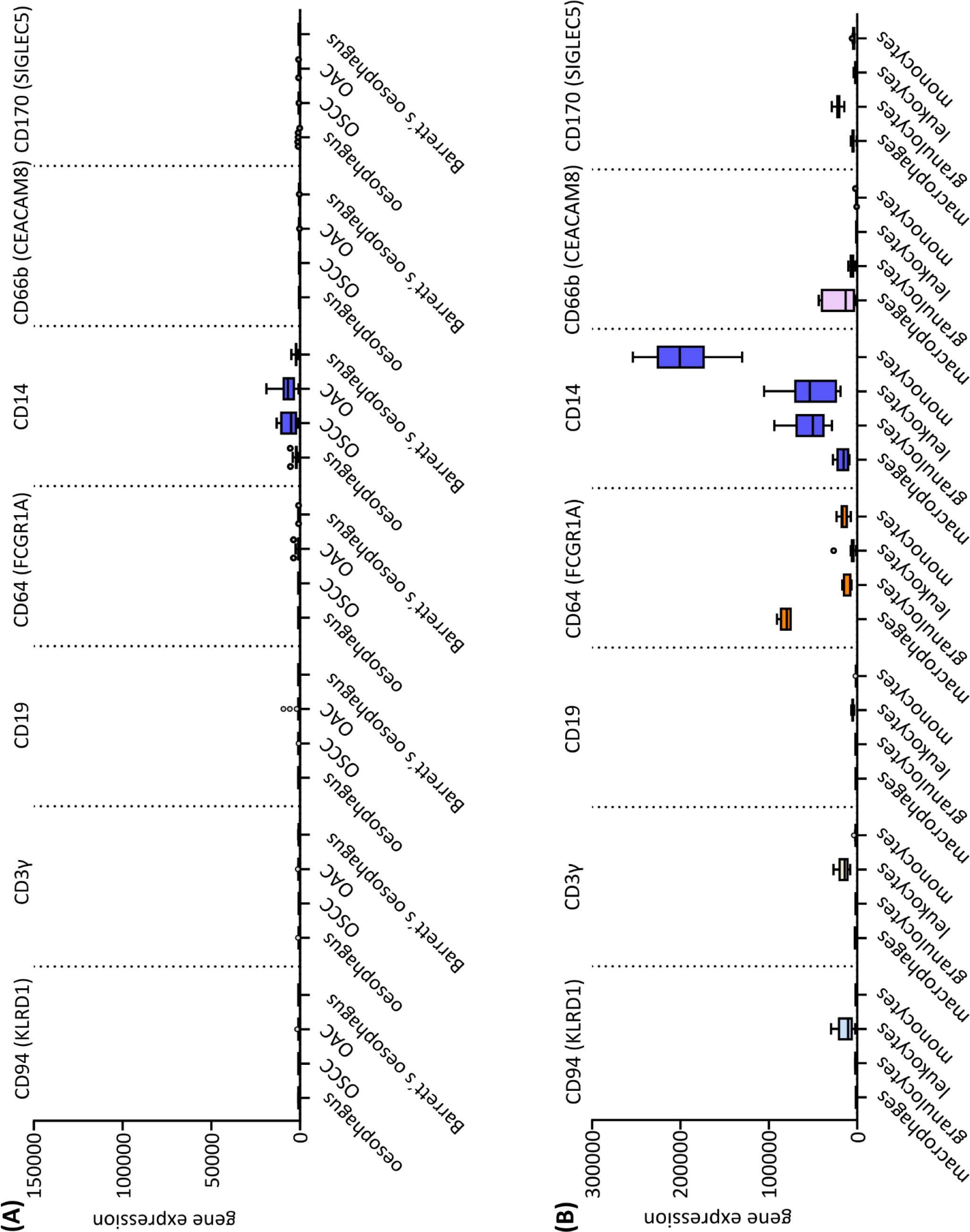
Gene expression levels of indicated immune cell markers: CD94 (NK-cells), CD3γ (T-cells), CD19 (B-cells), CD64 (monocytes/macrophages, Dendritic cells), CD14 (monocytes/macrophages, granulocytes), CD66b/CD67 (granulocytes), CD170 (tumour-associated neutrophiles, monocytes/macrophages, activated dendritic cells, lymphocytes-subsets) were analysed with the GENEVESTIGATOR platform **(A)** in oesophagus and cancerous oesophagus tissues (see Fig. 1G) and **(B)** in indicated human immune cell populations: macrophages (n = 6), granulocytes (n = 6), leukocytes (n = 8) and monocytes (n = 13).

**Figure S3:**
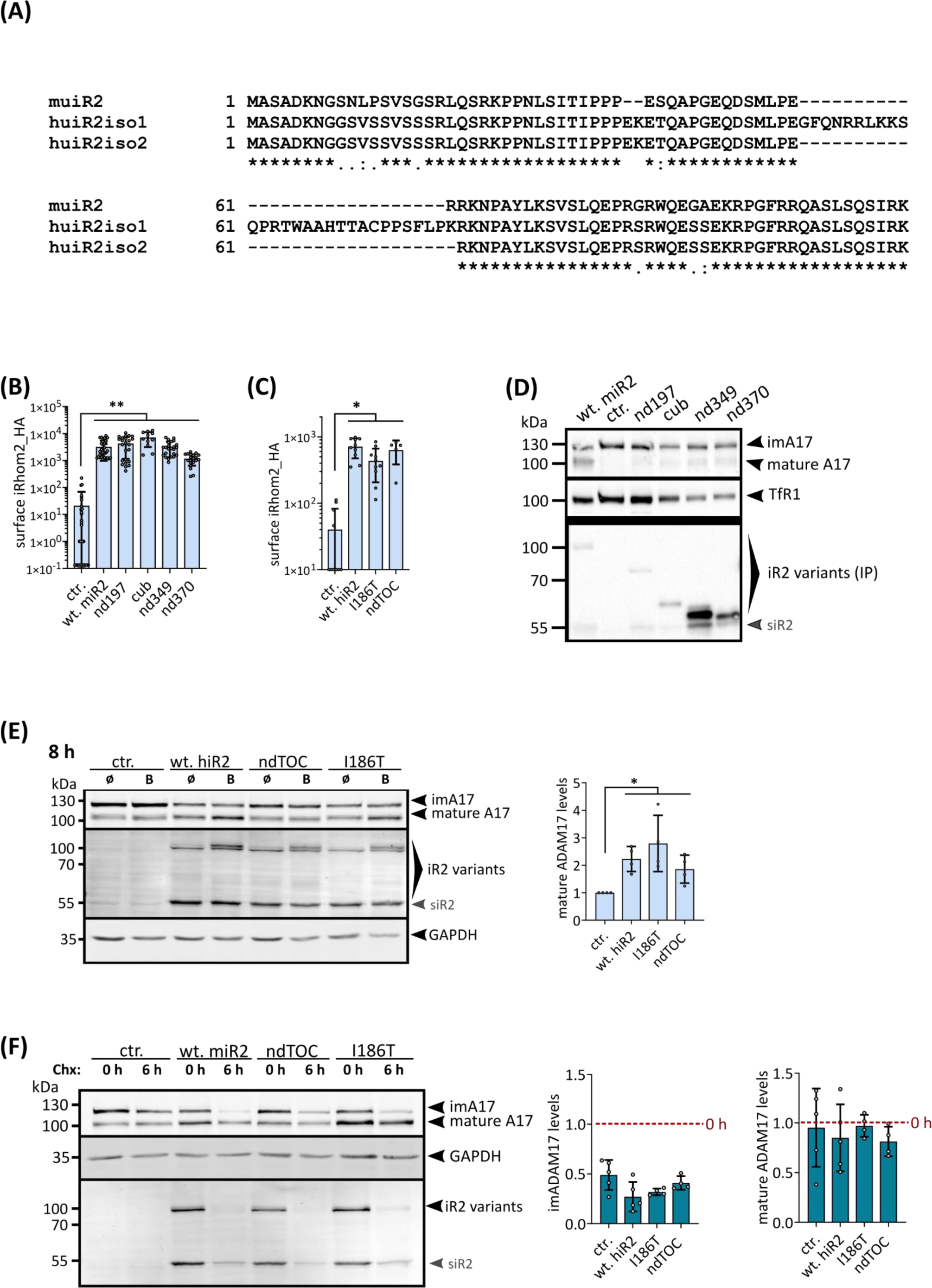
**(A)** Protein sequence alignment of the N-terminal region of the cytosolic tail of human iRhom2 (isoform 1 and isoform 2) and murine iRhom2. **(B)** Surface localisation of iRhom2 variants (with HA tag) in cells stably expressing the indicated murine constructs were measured by flow cytometry. For quantification, the geometric mean of the specific fluorescence signal was determined. n > 11. **(C)** Surface localisation of iRhom2 variants (with HA tag) in cells stably expressing the indicated human constructs were measured by flow cytometry. For quantification, the geometric mean of the specific fluorescence signal was determined. n > 5. **(D)** Immunoblot of samples from MEFs deficient for iRhom1 and iRhom2 stably expressing the indicated iRhom2 variants or GFP (ctr.) were used. The transferrin receptor (TfR1) served as an input control. (n = 3). **(E)** To analyse the fate of mature ADAM17 in cells expressing the indicated TOC mutations, cells were treated with bafilomycin (“B”) for 8 h to inhibit lysosomal degradation. Levels of mature ADAM17 were assessed by densitometric measurements and calculation of the ratio between mature ADAM17 and the loading control GAPDH, which is independent of lysosomal degradation. (n > 4). **(F)** To assess the turn-over rate of ADAM17 a cycloheximide-based (Chx) pulse-chase experiment was performed. Reduction of ADAM17 levels was assessed by densitometric measurements and normalisation to 0 h Chx treatment (set to 1). (n = 5).

**Figure S4:**
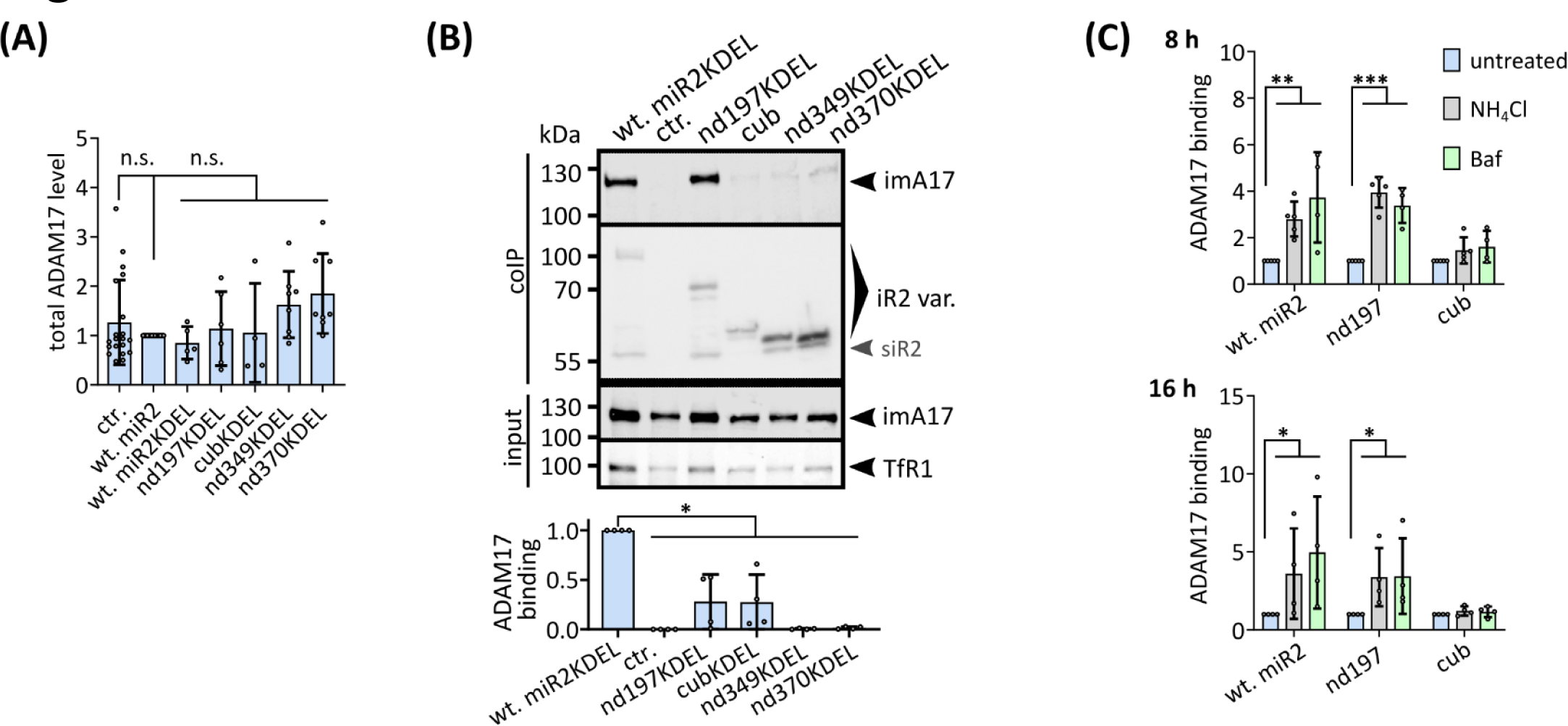
**(A)** ADAM17 levels of immunoblot (Fig. 6C) were assessed by densitometric measurements and calculation of the ratio between total ADAM17 (mature plus immature ADAM17) divided by the respective input control. Values were normalised to the respective wt-iRhom2 sample. n > 5. **(B-C)** To analyse interaction between ADAM17 and the indicated iRhom2 variants, coIPs were performed using the iRhom variants (with HA tag) as bait. Immunoblotting and subsequent densitometric measurements were used to quantitatively analyse iRhom2-ADAM17 binding (ratio of co-precipitated ADAM17 and precipitated iRhom2) and normalised to wt-iRhom2. **(B)** MEFs from mice deficient for iRhom1 and iRhom2 stably expressing indicated iRhom2 variants with ER retention signal KDEL or GFP (ctr.) were used (n = 4). **(C)** iRhom-ADAM17 binding after lysosomal inhibition was analysed. Densitometric measurements of coIP immunoblot (Fig. 6D) (n = 4).

